# Copper modulates iron-dependent survival through distinct TORC1 and AMPK signaling pathways

**DOI:** 10.64898/2026.05.18.725821

**Authors:** Arshia Naaz, Trishia Yi Ning Cheng, Jovian Jing Lin, Mingtong Gao, Rajkumar Dorajoo, Brian K. Kennedy, Mohammad Alfatah

## Abstract

Copper and iron are redox-active micronutrients with tightly coupled homeostasis, yet how copper modulates iron-dependent stress responses remains unclear. Using *Saccharomyces cerevisiae* under nutrient-limited conditions, we uncoupled proliferative growth from long-term survival to dissect metal-dependent adaptation. Copper selectively preserved survival without affecting growth, whereas iron showed similar effects. Iron chelation impaired growth and suppressed electron transport chain gene expression; copper partially rescued these defects but required iron for its pro-survival activity. Despite this interdependence, copper and iron engaged distinct signaling programs. Iron-dependent survival required a Target of Rapamycin complex 1 (TORC1)-permissive state and was attenuated by rapamycin, whereas copper remained active under TORC1 inhibition. In contrast, copper promoted survival through AMP-activated protein kinase (AMPK) and antioxidant pathways, while iron exhibited context-dependent AMPK reliance. Together, these findings identify copper and iron as state-dependent regulators of cellular survival.

## INTRODUCTION

Copper and iron are essential redox-active micronutrients that play central roles in cellular metabolism, mitochondrial function, and stress adaptation ^1–13^. Their ability to cycle between oxidation states enables key biochemical processes, including respiration and antioxidant defense, but also poses a risk of oxidative damage if not tightly controlled ^14–18^. Accordingly, copper and iron homeostasis are closely coordinated, and perturbation of one metal can influence the availability and function of the other ^10,19–24^.

This interdependence is particularly evident in pathways linking metal availability to metabolic and redox processes. Iron is required for heme and iron–sulfur cluster biogenesis, whereas copper supports enzymes involved in oxidative metabolism and antioxidant defense^3,6,9–11,13,20,25–27^. Disruption of either metal alters redox balance and cellular fitness^4,6,8,11,22,24,28–34^. However, despite extensive characterization of their individual roles, how copper availability modulates iron-dependent survival and whether these micronutrients engage distinct regulatory pathways remain incompletely understood.

Cells frequently encounter nutrient and environmental limitations in physiological contexts. Aging tissues, ischemic regions, and solid tumors often experience restricted nutrient and oxygen availability, imposing metabolic and redox stress that necessitates adaptive survival programs ^35–42^. Under these conditions, cells undergo transitions from proliferative growth to maintenance states, requiring coordinated regulation of nutrient sensing, mitochondrial function, and stress-response pathways.

Cellular responses to metabolic stress are governed by conserved signaling networks, including Target of Rapamycin Complex 1 (TORC1) and AMP-activated protein kinase (AMPK) ^43–62^. TORC1 promotes anabolic growth under nutrient-replete conditions, whereas AMPK (Snf1 in yeast) supports adaptation to energetic and oxidative stress. While these pathways have been extensively studied, how redox-active micronutrients such as copper and iron interface with these signaling systems to regulate survival across distinct metabolic states remains poorly defined.

The budding yeast *Saccharomyces cerevisiae* provides a powerful model to dissect these relationships, as it enables separation of proliferative growth from long-term stationary-phase survival under defined nutrient conditions ^63–66^. This system allows systematic interrogation of how metabolic state, redox balance, and signaling pathways integrate to determine cellular fate.

Here, we investigate how copper availability modulates iron-dependent metabolic and redox adaptation across distinct cellular states. Using dose–response analyses, iron chelation, and genetic and pharmacological perturbations of TORC1 and AMPK/Snf1 signaling, we show that copper and iron promote survival through mechanistically distinct pathways. We demonstrate that iron-dependent survival requires a TORC1-permissive metabolic state and is associated with sustained TORC1 activity, whereas copper retains pro-survival function under TORC1 limitation and engages stress-adaptive pathways. Together, these findings reveal that redox-active micronutrients regulate cellular state transitions between growth and survival through coordinated integration of nutrient signaling and stress-response networks.

## RESULTS

### Copper selectively preserves survival without affecting proliferative growth under nutrient limitation

To determine how copper availability influences cellular proliferation and long-term survival under nutrient-limited conditions, we used the prototrophic Saccharomyces cerevisiae strain CEN.PK grown in synthetic defined (SD) medium.

Mitotic growth was monitored by OD600 at 24, 48, and 72 h following supplementation with copper (II) sulfate (CuSO_4_) across a 12-point concentration range (0–200 µM; 2-fold serial dilution). Copper supplementation did not alter growth kinetics compared with untreated controls across all tested concentrations **(Figure 1A**), indicating that proliferative capacity is preserved within this range.

**Figure 1.**
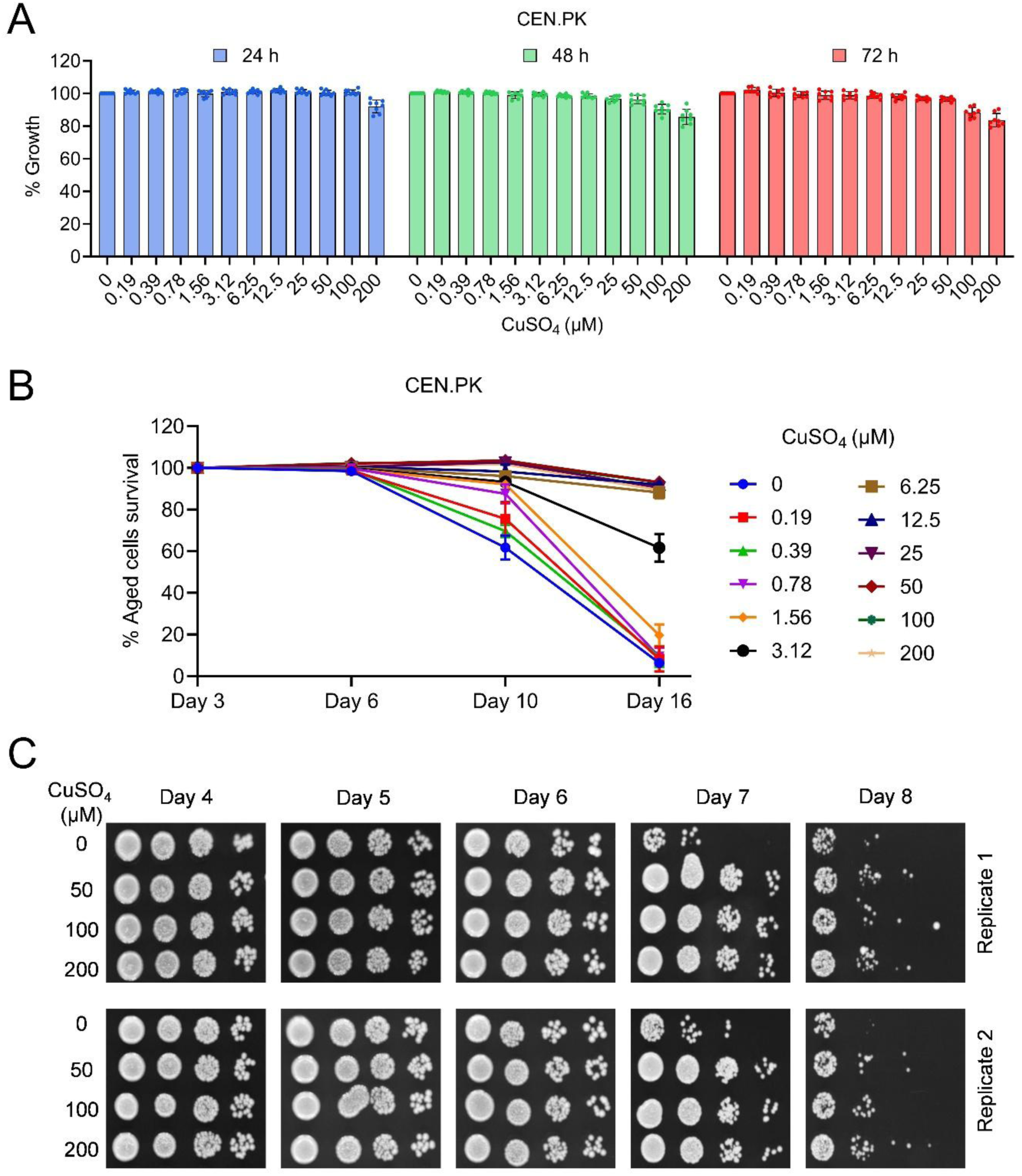
Copper preserves long-term survival without affecting proliferative growth. (A) Growth of Saccharomyces cerevisiae CEN.PK cells cultured in synthetic defined (SD) medium supplemented with increasing concentrations of CuSO_4_ (0–200 µM). Cell density was measured at 24, 48, and 72 h by OD600 and normalized to untreated controls. Data represent mean ± SD (n = 8). (B) Chronological survival of CEN.PK cells cultured in the presence of CuSO_4_. Viability was assessed at indicated time points (days 3, 6, 10, and 16) using a liquid outgrowth assay and expressed relative to day 3. Data represent mean ± SD (n = 4). (C) Representative spot dilution assay of aged cultures grown with CuSO_4_ (0, 50, 100, and 200 µM). Serial dilutions were plated on YPD agar at indicated time points (days 4–8). Two independent biological replicates are shown.

We next assessed long-term survival using two independent outgrowth-based assays. In the liquid outgrowth assay, stationary-phase cultures were sampled at indicated time points and reinoculated into fresh YPD medium in 96-well format, where regrowth served as a quantitative proxy for viability. Copper supplementation preserved long-term survival in a concentration-dependent manner. By day 16, viability reached ∼60% at 3.12 µM CuSO₄ and approached ∼100% across 6.25–200 µM, compared with <10% in untreated cultures **(Figure 1B)**.

Consistent with these findings, a spotting-based recovery assay showed markedly enhanced colony outgrowth from copper-treated cultures relative to untreated controls **(Figure 1C)**, confirming improved recovery following prolonged nutrient limitation.

To assess the generality of this phenotype, we performed parallel experiments in the auxotrophic strain BY4743 grown in supplemented SD medium. Consistent with the CEN.PK strain, copper preserved long-term survival without affecting growth in BY4743 cells **(Figures S1A and S1B).**

These findings demonstrate that copper selectively promotes a survival-associated cellular state without altering proliferative growth under nutrient-limited conditions.

### Copper and iron exhibit convergent pro-survival effects, with iron acting at lower effective doses

Copper and iron homeostasis are tightly interdependent across eukaryotes ^3–13^. Building on our previous observations and other studies showing that iron supplementation enhances stationary-phase survival ^67–70^, we directly compared copper and iron dose–response profiles under identical conditions. CEN.PK cells were supplemented with copper (II) sulfate (CuSO_4_) or iron (II) sulfate (FeSO_4_) across a 12-point concentration range (0–200 µM; 2-fold serial dilution). Neither CuSO_4_ or FeSO_4_ altered growth kinetics across the tested range **(Figures S2A and S2B)**, indicating that both metals are well tolerated during proliferative conditions.

We next quantified stationary-phase viability using the liquid outgrowth assay. Consistent with Figure 1B, CuSO_4_ at 6.25–200 µM preserved near-complete viability by day 16 **(Figure 2A)**. FeSO_4_ produced a similarly robust, and in some cases stronger, pro-survival effect, maintaining near-complete viability across a broader concentration range (∼0.39–200 µM) **(Figure 2B)**. These results indicate that iron promotes survival at lower effective concentrations than copper, suggesting greater potency under these conditions.

**Figure 2.**
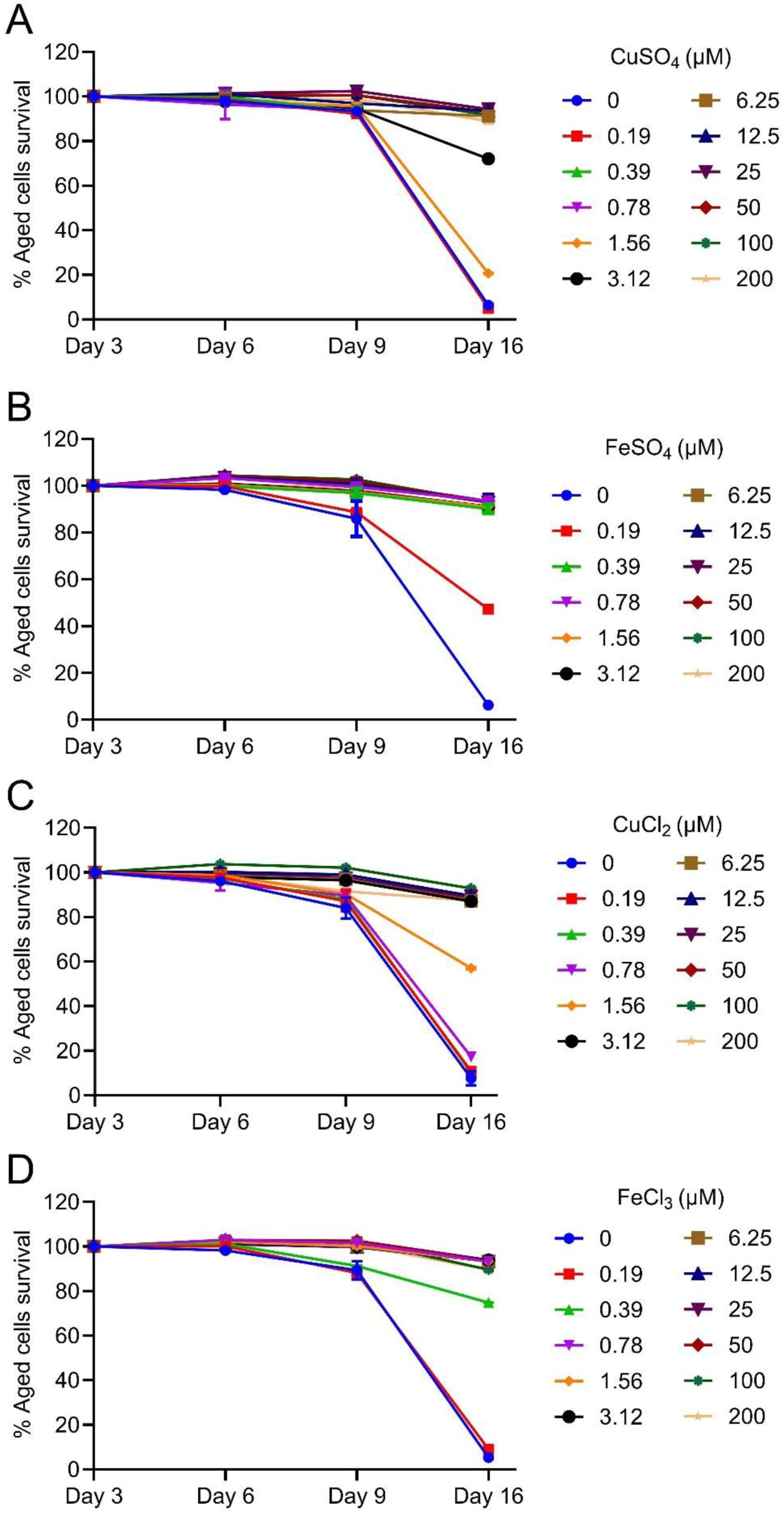
Copper and iron exhibit convergent pro-survival effects. (A) Chronological survival of *Saccharomyces cerevisiae* CEN.PK cells cultured in synthetic defined (SD) medium supplemented with increasing concentrations of CuSO_4_ (0–200 µM). Viability was assessed at indicated time points (days 3, 6, 9, and 16) using a liquid outgrowth assay and expressed relative to day 3. Data represent mean ± SD (n = 2). (B) Chronological survival of CEN.PK cells cultured in the presence of FeSO_4_ (0–200 µM), assessed as in (A). Data represent mean ± SD (n = 2). (C) Chronological survival of CEN.PK cells cultured in the presence of CuCl_2_ (0–200 µM), assessed as in (A). Data represent mean ± SD (n = 2). (D) Chronological survival of CEN.PK cells cultured in the presence of FeCl_3_ (0–200 µM), assessed as in (A). Data represent mean ± SD (n = 2).

To determine whether these effects depend on the counterion, we evaluated copper (II) chloride (CuCl_2_) and iron (III) chloride (FeCl_3_). Both salts recapitulated the central trends observed with sulfate forms, showing no detectable impact on growth and robust preservation of stationary-phase viability in a dose-dependent manner **(Figures 2C, 2D and S2C–S2D)**.

These findings demonstrate that copper and iron converge on survival regulation but differ in potency and effective dose range, consistent with shared roles in metabolic and redox processes.

### Iron chelation suppresses growth and downregulates mitochondrial electron transport chain gene programs

To determine whether copper-dependent survival is linked to iron metabolism, we induced functional iron limitation using bathophenanthrolinedisulfonic acid (BPS), a ferrous iron (Fe²⁺) chelator ^71^. Cells were cultured in SD medium containing BPS, with or without supplemental iron, and mitotic growth was monitored. BPS caused a pronounced growth defect, consistent with the essential role of iron in cellular proliferation, while iron supplementation restored growth toward control levels **(Figures 3A, 3B and S3A)**, confirming that the phenotype arises from iron limitation.

**Figure 3.**
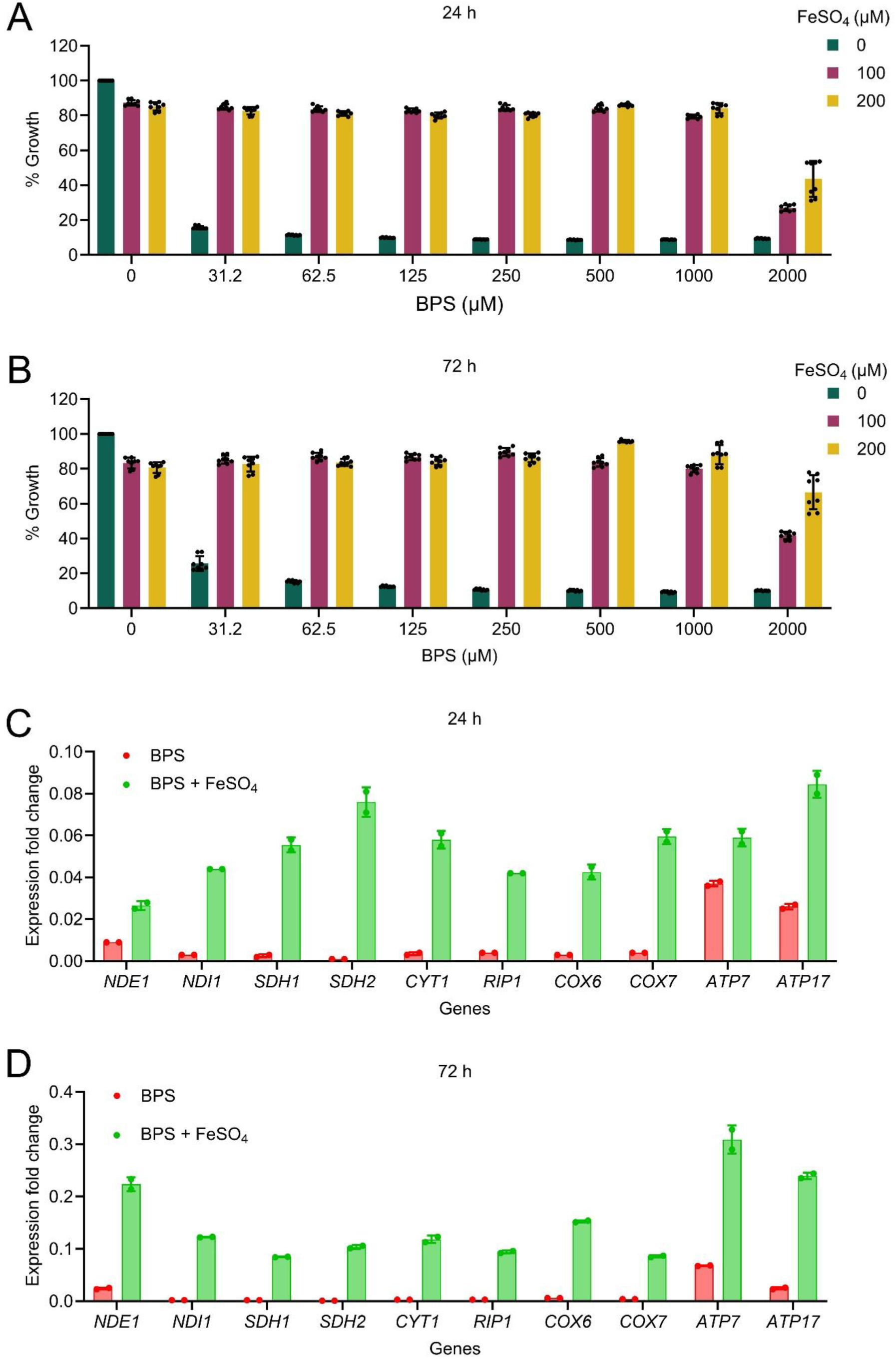
Iron supplementation rescues growth and restores mitochondrial gene expression under iron chelation. (A) Growth of *Saccharomyces cerevisiae* CEN.PK cells cultured in synthetic defined (SD) medium with increasing concentrations of the iron chelator BPS (0–2000 µM) in the presence or absence of FeSO_4_ (0, 100, and 200 µM). Cell density was measured at 24 h by OD600 and normalized to untreated controls. (B) Growth of CEN.PK cells under the same conditions as in (A), measured at 72 h. (C) Relative expression of mitochondrial electron transport chain–related genes (*NDE1, NDI1, SDH1, SDH2, CYT1, RIP1, COX6, COX7, ATP7*, and *ATP17*) in cells treated with BPS alone or BPS supplemented with FeSO_4_. Gene expression was measured at 24 h by quantitative RT–PCR, normalized to ACT1, and expressed relative to untreated control (Figure S3B). (D) Relative gene expression as in (C), measured at 72 h. (Figure S3C).

Given the central role of iron in heme and iron–sulfur cluster biogenesis, we next assessed whether iron chelation alters mitochondrial respiratory programs. For gene expression analysis, cells were grown in larger-volume flask cultures to enable sufficient biomass collection for RNA extraction. We quantified representative genes spanning all major electron transport chain (ETC) complexes: *NDE1/NDI1* (Complex I), *SDH1/SDH2* (Complex II), *CYT1/RIP1* (Complex III), *COX6/COX7* (Complex IV), and *ATP7/ATP17* (Complex V) ^67,72^. Iron chelation significantly reduced the expression of these ETC-associated genes, while iron supplementation restored their expression **(Figures 3C, 3D, S3B and S3C)**. Notably, iron supplementation also rescued the expression of tricarboxylic acid (TCA) cycle genes **(Figures S3D and S3E)**.

Taken together, these findings demonstrate that iron limitation suppresses mitochondrial respiratory gene programs, establishing a state of impaired mitochondrial and redox capacity. This provides a mechanistic framework to assess how copper modulates survival under iron-dependent mitochondrial stress.

### Iron availability is required for copper-dependent survival benefits

Having established that BPS induces functional iron limitation and suppresses mitochondrial gene programs, we next asked whether copper can compensate for iron chelation–associated defects. Across multiple growth assays, copper supplementation partially rescued BPS-induced growth impairment in a concentration-dependent manner **(Figures 4A–4C and S4A–S4C)**, indicating that copper can support proliferative capacity under iron-limited conditions. This is consistent with partially overlapping or coordinated roles of copper and iron in core metabolic processes ^3,6,9,10,13,19,21^.

**Figure 4.**
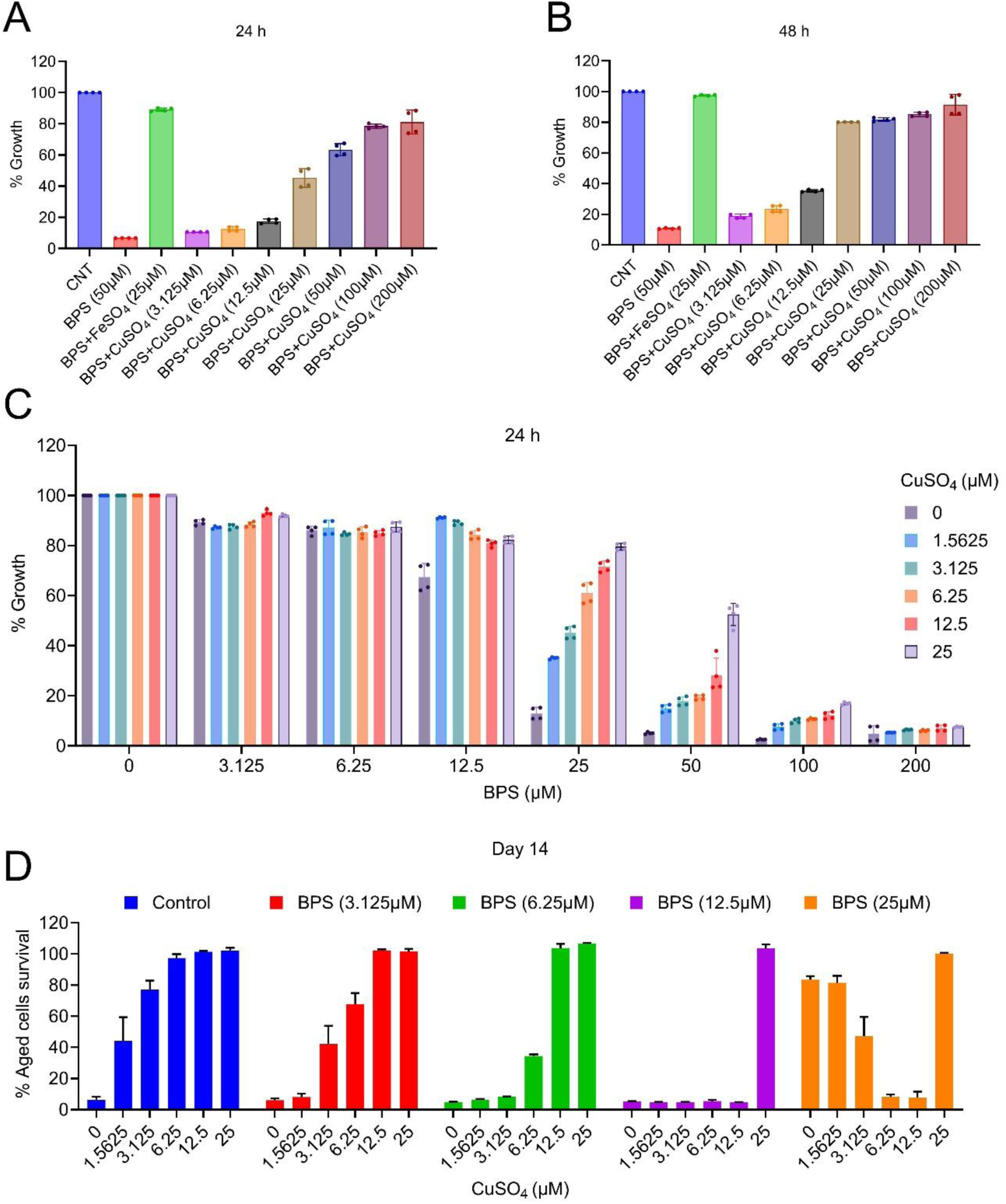
Copper partially rescues growth and survival under iron chelation. (A) Growth of *Saccharomyces cerevisiae* CEN.PK cells cultured in synthetic defined (SD) medium under control (CNT), BPS (50 µM), BPS + FeSO_4_ (25 µM), or BPS supplemented with increasing concentrations of CuSO_4_ (3.125–200 µM). Cell density was measured at 24 h by OD600 and normalized to control. Data represent mean ± SD (n = 4). (B) Growth of CEN.PK cells under the same conditions as in (A), measured at 48 h. Data represent mean ± SD (n = 4). (C) Growth of CEN.PK cells cultured with increasing concentrations of BPS (0–200 µM) in the presence of CuSO_4_ (0–25 µM). Cell density was measured at 24 h by OD600 and normalized to untreated controls. Data represent mean ± SD (n = 4). (D) Chronological survival of CEN.PK cells cultured with increasing concentrations of CuSO_4_ (0–25 µM) in the presence of BPS (3.125–25 µM). Viability was assessed at day 14 using a liquid outgrowth assay and expressed relative to day 3.

We then tested whether copper-dependent preservation of stationary-phase viability requires iron availability. Stationary-phase survival was quantified across increasing concentrations of BPS in the presence of copper. Progressive iron chelation attenuated the pro-survival effects of copper in a dose-dependent manner. At 6.25 µM BPS, the survival benefit conferred by low copper concentrations (1.56–3.125 µM) was eliminated **(Figure 4D)**. At higher copper levels (6.25 µM), BPS partially suppressed the effect, reducing day 16 viability from near-complete survival to ∼40%. At 12.5 µM BPS, copper-dependent survival benefits across all tested copper concentrations (1.56–12.5 µM) were abolished **(Figure 4D)**.

Together, these findings indicate that copper-mediated survival requires iron availability, supporting a model in which copper acts through iron-dependent metabolic and redox processes rather than functioning independently. This establishes iron as a necessary component of copper-driven survival adaptation under stress conditions.

### Severe iron chelation induces a distinct survival state that is antagonized by low copper

To investigate the effects of more severe iron limitation, we examined 25 µM BPS. Unexpectedly, this condition alone preserved stationary-phase viability even in the absence of copper **(Figures 4D, S4D and S4E)**, indicating that strong iron chelation can induce a distinct adaptive survival state.

Growth analysis revealed that 25 µM BPS delayed proliferation at 24 h (∼20% of control) but partially recovered by 72 h, reaching ∼80% of control **(Figures S4A–S4D)**, corresponding to the onset of stationary-phase measurements. This slow-growth phenotype is consistent with metabolic reprogramming associated with reduced nutrient signaling, including TORC1 suppression ^59,73–75^. In line with this, iron limitation has been linked to modulation of TORC1 activity ^76–83^, suggesting that severe iron depletion may promote survival through coordinated changes in metabolic and signaling state.

Strikingly, the survival advantage conferred by 25 µM BPS was antagonized by co-treatment with low-to-moderate copper concentrations (1.56–12.5 µM), which reduced viability relative to BPS alone **(Figure 4D)**. This indicates that, under conditions of strong iron limitation, low copper disrupts an iron-deficiency–induced adaptive survival program. In contrast, higher copper levels (25 µM) retained pro-survival activity in the presence of 6.25–12.5 µM BPS and did not diminish the survival benefit associated with severe iron chelation **(Figure 4D)**.

Together, these findings reveal a non-linear interaction between copper and iron in which copper can either support or interfere with survival depending on the underlying metabolic state. This suggests that copper modulates cellular adaptation through iron-dependent metabolic and redox processes, while also influencing broader state transitions under chronological stress.

### Copper and iron exhibit distinct interactions with pharmacological TORC1 inhibition

Given that severe iron chelation phenocopies aspects of nutrient signaling ^59,73,74,76–83^, we examined how copper and iron interact with pharmacological inhibition of TORC1 using rapamycin. Both iron (5–10 µM) and copper (5–10 µM) preserved long-term survival in wild-type cells. Low-dose rapamycin (0.31–2.5 nM) alone had minimal effects but attenuated the survival benefits of metal supplementation, indicating functional interplay between metal-dependent and TORC1-regulated survival pathways **(Figures 5A and 5B).**

**Figure 5.**
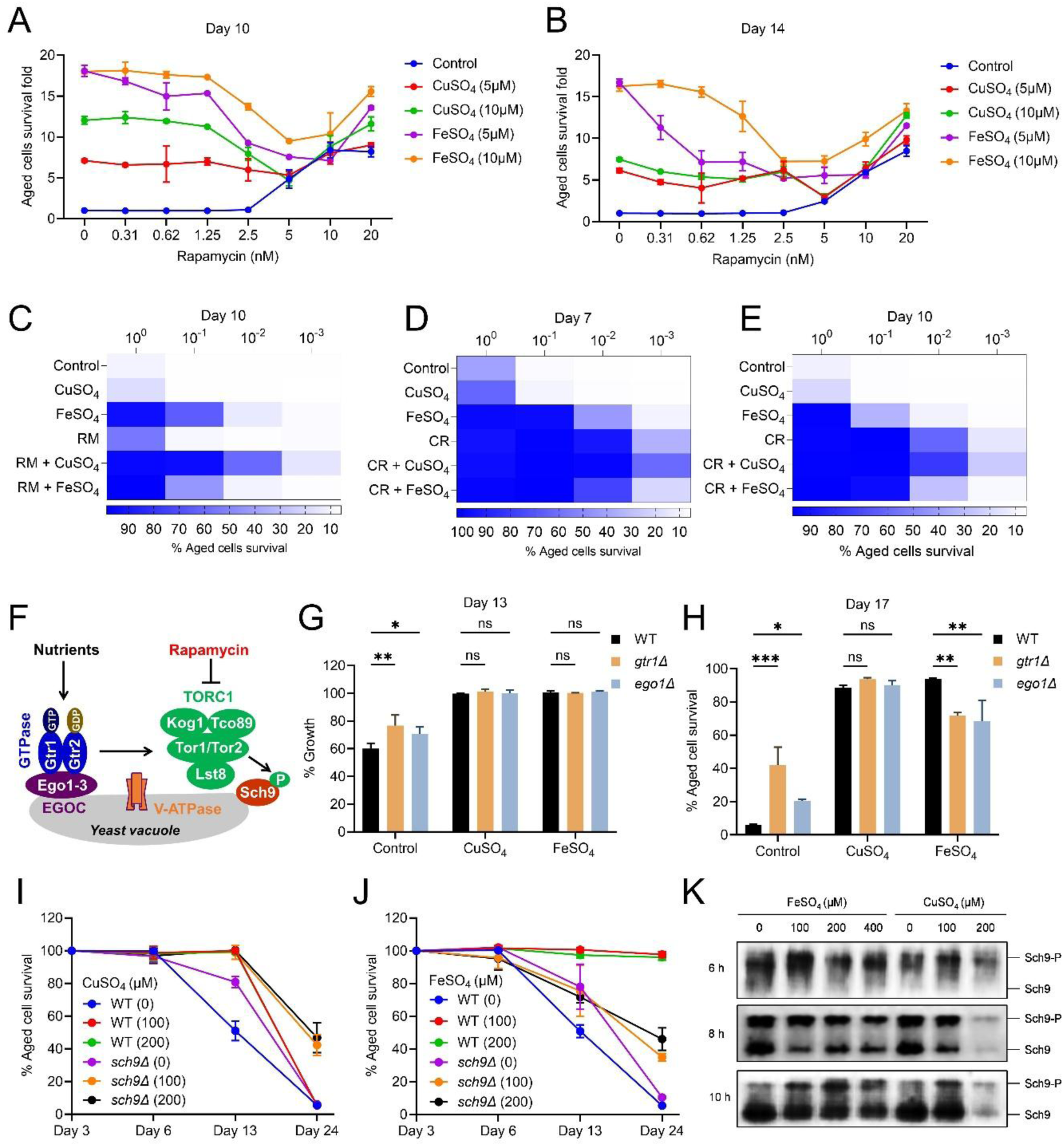
Copper and iron differentially interact with TORC1 pathway. (A–B) Chronological survival of *Saccharomyces cerevisiae* CEN.PK cells cultured with increasing concentrations of rapamycin (0–20 nM) in the presence or absence of CuSO_4_ (5 and 10 µM) or FeSO_4_ (5 and 10 µM). Viability was assessed at day 10 (A) and day 14 (B) using a liquid outgrowth assay and expressed relative to day 3. Data represent mean ± SD (n = 2). (C) Heatmap representation of chronological survival of CEN.PK cells cultured under control conditions, CuSO_4_, FeSO_4_, rapamycin (RM), and their combinations at day 10. Serial 10-fold dilutions were used for outgrowth-based quantification. Data represent mean ± SD (n = 2). (D–E) Heatmap representation of chronological survival under calorie restriction (CR) conditions and in combination with CuSO_4_ or FeSO_4_. Viability was assessed at day 7 (D) and day 10 (E). Data represent mean ± SD (n = 2). (F) Schematic representation of TORC1 signaling components and regulatory modules in yeast. (G) Growth of wild-type (WT), *gtr1Δ*, and *ego1Δ* strains cultured under control conditions or supplemented with CuSO_4_ or FeSO_4_. Cell density was measured by OD600 and normalized to WT control. Data represent mean ± SD (n = 2). (H) Chronological survival of WT, *gtr1Δ*, and *ego1Δ* strains cultured under the indicated conditions. Viability was assessed at day 17 using a liquid outgrowth assay. Data represent mean ± SD (n = 2). (I–J) Chronological survival of WT and *sch9Δ* strains cultured with CuSO_4_ (I) or FeSO_4_ (J) at indicated concentrations. Viability was assessed at indicated time points and expressed relative to day 3. Data represent mean ± SD (n = 2). (K) Immunoblot analysis of Sch9 phosphorylation (Sch9-P) and total Sch9 levels in cells treated with CuSO_4_ or FeSO_4_ at indicated concentrations. Samples were collected at 6 h, 8 h, and 10 h.

Notably, the nature of this interaction differed between the two metals. Iron exhibited a consistently antagonistic relationship with rapamycin, with its pro-survival effect markedly reduced even at higher rapamycin concentrations. In contrast, copper displayed a dose-dependent interaction: while modest at low rapamycin levels, copper enhanced survival at higher rapamycin concentrations (10–20 nM), consistent with a cooperative effect. These findings indicate that iron remains incompatible with TORC1 inhibition, whereas copper becomes beneficial under stronger TORC1 suppression **(Figures 5A, 5B and S5A–S5C).**

These observations were confirmed using dilution-based outgrowth assays, which showed that copper co-treatment enhanced survival in the presence of rapamycin, whereas iron co-treatment reduced survival relative to iron alone **(Figure 5C).**

Together, these results reveal opposing pharmacological interactions of copper and iron with TORC1 inhibition.

### Copper and iron differentially modulate survival under calorie restriction

We next examined whether these interactions extend to calorie restriction (CR), a physiological condition associated with reduced TORC1 activity ^84–91^. As expected, glucose restriction (0.5%) increased long-term survival compared with nutrient-replete conditions **(Figures 5D and 5E).**

Under CR conditions, copper supplementation further enhanced survival, indicating a cooperative interaction with reduced nutrient signaling. In contrast, iron supplementation attenuated the survival benefit conferred by CR, demonstrating an antagonistic effect in this context **(Figures 5D and 5E).**

These findings parallel the effects observed with pharmacological TORC1 inhibition and indicate that copper and iron differentially influence survival under both chemical and physiological suppression of TORC1 signaling.

Together, these results extend the divergence between copper and iron to physiological TORC1 modulation.

### Iron-dependent survival requires TORC1 signaling, whereas copper acts independently

To determine whether these effects depend on TORC1 signaling, we examined genetic perturbations of the pathway **(Figure 5F)** ^60,92–97^. Deletion of the upstream TORC1 activators *GTR1* and *EGO1* increased baseline survival relative to wild-type cells, consistent with reduced TORC1 activity **(Figures 5G, 5H, S5D and S5E)**. In these backgrounds, iron supplementation conferred a diminished survival benefit compared with wild-type cells, indicating that iron-mediated survival requires intact upstream TORC1 signaling. In contrast, copper maintained its pro-survival effect in both *gtr1Δ* and *ego1Δ* strains, suggesting reduced dependence on upstream TORC1 activation.

We next examined downstream TORC1 signaling using deletion of SCH9, a major effector of TORC1 **(Figure 5F)** ^90,98–100^. As expected, *sch9Δ* cells exhibited elevated baseline survival **(Figures 5I, 5J, S5F and S5G)**. In this background, iron supplementation reduced survival relative to *sch9Δ* alone, further supporting that iron-dependent survival requires TORC1–Sch9 signaling. In contrast, copper supplementation further enhanced survival in *sch9Δ* cells, indicating that its pro-survival effects are at least partially independent of TORC1 output.

Consistent with these genetic findings, analysis of TORC1 activity revealed distinct biochemical responses. Sch9 phosphorylation decreased during the transition from exponential to early stationary phase in untreated cells, reflecting reduced TORC1 activity upon growth arrest **(Figure 5K)**. Iron supplementation maintained elevated Sch9 phosphorylation across this transition, indicating sustained TORC1 activity under nutrient-limited conditions. In contrast, copper had only a modest effect and did not prevent the decline in TORC1 activity.

Together, these findings establish that iron-dependent survival is tightly coupled to TORC1 signaling, whereas copper promotes survival through a partially TORC1-independent mechanism.

### Copper promotes stress survival through AMPK/Snf1-dependent metabolic adaptation

Given copper’s activity under TORC1-limited conditions, we next examined whether copper supports survival under metabolic stress. AMPK is a conserved regulator of energy homeostasis and stress adaptation, with Snf1 serving as its yeast ortholog ^45–54,59,60,62^. As expected, *snf1Δ* cells exhibited reduced stationary-phase survival compared with wild-type controls **(Figure 6A)**, consistent with impaired stress adaptation ^67,101^.

**Figure 6.**
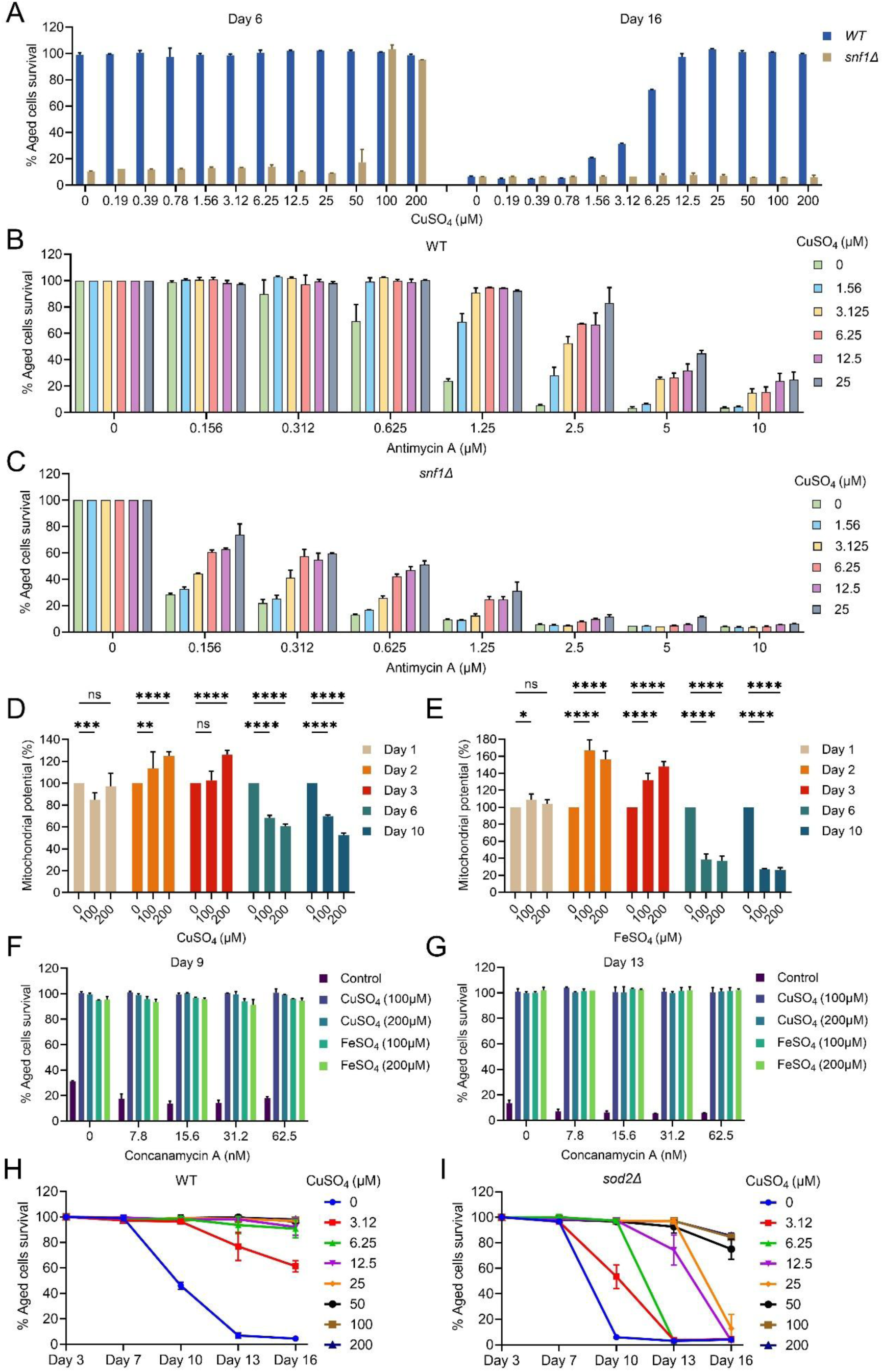
Copper supports survival under metabolic and oxidative stress through Snf1/AMPK-associated pathways. (A) Chronological survival of wild-type (WT) and *snf1Δ* strains cultured in synthetic defined (SD) medium supplemented with increasing concentrations of CuSO_4_ (0–200 µM). Viability was assessed at day 6 and day 16 using a liquid outgrowth assay and expressed relative to day 2. Data represent mean ± SD (n = 2). (B–C) Chronological survival of WT (B) and *snf1Δ* (C) cells cultured with increasing concentrations of antimycin A (0–10 µM) in the presence of CuSO_4_ (0–25 µM). Viability was assessed using a liquid outgrowth assay and expressed relative to day 2. Data represent mean ± SD (n = 2). (D–E) Mitochondrial membrane potential in WT cells treated with CuSO_4_ (0–200 µM) (D) or FeSO_4_ (0–200 µM) (E) at indicated time points (days 1, 2, 3, 6, and 10). Fluorescence intensity was measured and normalized to untreated controls. Data represent mean ± SD (n = 4). (F–G) Chronological survival of CEN.PK cells cultured with concanamycin A (0–62.5 nM) in the presence or absence of CuSO_4_ or FeSO_4_ (100 and 200 µM). Viability was assessed at day 9 (F) and day 13 (G). Data represent mean ± SD (n = 2). (H–I) Chronological survival of WT (H) and *sod2Δ* (I) strains cultured with increasing concentrations of CuSO_4_ (0–200 µM). Viability was assessed at indicated time points and expressed relative to day 3. Data represent mean ± SD (n = 2).

Copper-dependent survival benefits were attenuated in *snf1Δ* cells and required higher concentrations to achieve partial rescue, indicating that AMPK/Snf1 contributes to copper-mediated survival **(Figure 6A)**. To further assess this relationship under respiratory stress, we used antimycin A (AMA), a complex III inhibitor ^102^. AMA reduced survival in wild-type cells and caused increased sensitivity in *snf1Δ* cells **(Figures S6A and S6B)**, consistent with a requirement for Snf1 in stress adaptation. Copper supplementation improved survival in AMA-treated wild-type cells in a dose-dependent manner, whereas this protective effect was diminished in *snf1Δ* cells **(Figures 6B, S6C and S6D)**, supporting a Snf1-dependent mechanism.

Consistent with these findings, copper treatment increased mitochondrial membrane potential at early time points (day 2–3), followed by a decline at later stages (day 6–10), indicating a transient enhancement of mitochondrial activity during stress adaptation **(Figure 6D)**. Iron also increased mitochondrial membrane potential at early time points, with a greater magnitude than copper **(Figure 6E)**. However, this response was less sustained, with a sharper decline at later stages. These data suggest that both metals promote early mitochondrial activation, but copper induces a more moderate and temporally stable response, whereas iron elicits a stronger but less durable effect.

To assess organelle-level stress, we examined inhibition of the vacuolar H⁺-ATPase using concanamycin A ^69,103–105^. Disruption of vacuolar function reduced stationary-phase survival, consistent with its role in mitochondrial homeostasis and aging ^69^. Notably, both copper and iron supplementation restored survival under these conditions **(Figures 6F and 6G)**, indicating that metal availability can buffer organelle dysfunction and support survival under metabolic stress.

Together, these findings indicate that copper promotes stress survival through Snf1/AMPK-dependent metabolic adaptation, accompanied by controlled modulation of mitochondrial function.

### Copper and iron differentially engage antioxidant defense pathways

We next investigated whether antioxidant capacity contributes to metal-dependent survival. *SOD2*, encoding a mitochondrial superoxide dismutase, is essential for limiting oxidative damage under respiratory conditions ^63,106,107^. As expected, *sod2Δ* cells exhibited reduced stationary-phase survival **(Figures 6H and 6I)**.

Copper-dependent survival benefits were diminished in *sod2Δ* cells and required higher concentrations for partial rescue **(Figure 6I and S6E)**, indicating that antioxidant defense contributes to copper-mediated survival. In contrast, iron supplementation robustly restored survival in *sod2Δ* cells even at low concentrations **(Figures S6F–S6H)**, suggesting that iron-mediated survival occurs largely independently of *SOD2* and may instead rely on alternative mechanisms that support mitochondrial function.

These results reveal a key mechanistic distinction: copper-dependent survival requires functional antioxidant pathways, whereas iron can promote survival even under compromised redox defense.

Together, these findings demonstrate that copper and iron differentially engage redox adaptation, with copper relying on antioxidant defense and iron acting through alternative mitochondrial-supporting mechanisms.

### AMPK/Snf1 modulates copper- and iron-dependent oxidative stress tolerance

To determine how metal supplementation influences redox adaptation, we assessed oxidative stress tolerance using hydrogen peroxide (H_2_O_2_). Both wild-type and *snf1Δ* cells exhibited sensitivity to H_2_O_2_-induced cytotoxicity across 24, 48, and 72 h time points **(Figures 7A and 7B)**. However, *snf1Δ* cells were consistently more sensitive, particularly as cultures approached stationary phase, indicating that Snf1 is required for effective stress adaptation under these conditions, consistent with previous studies ^67,101^.

**Figure 7.**
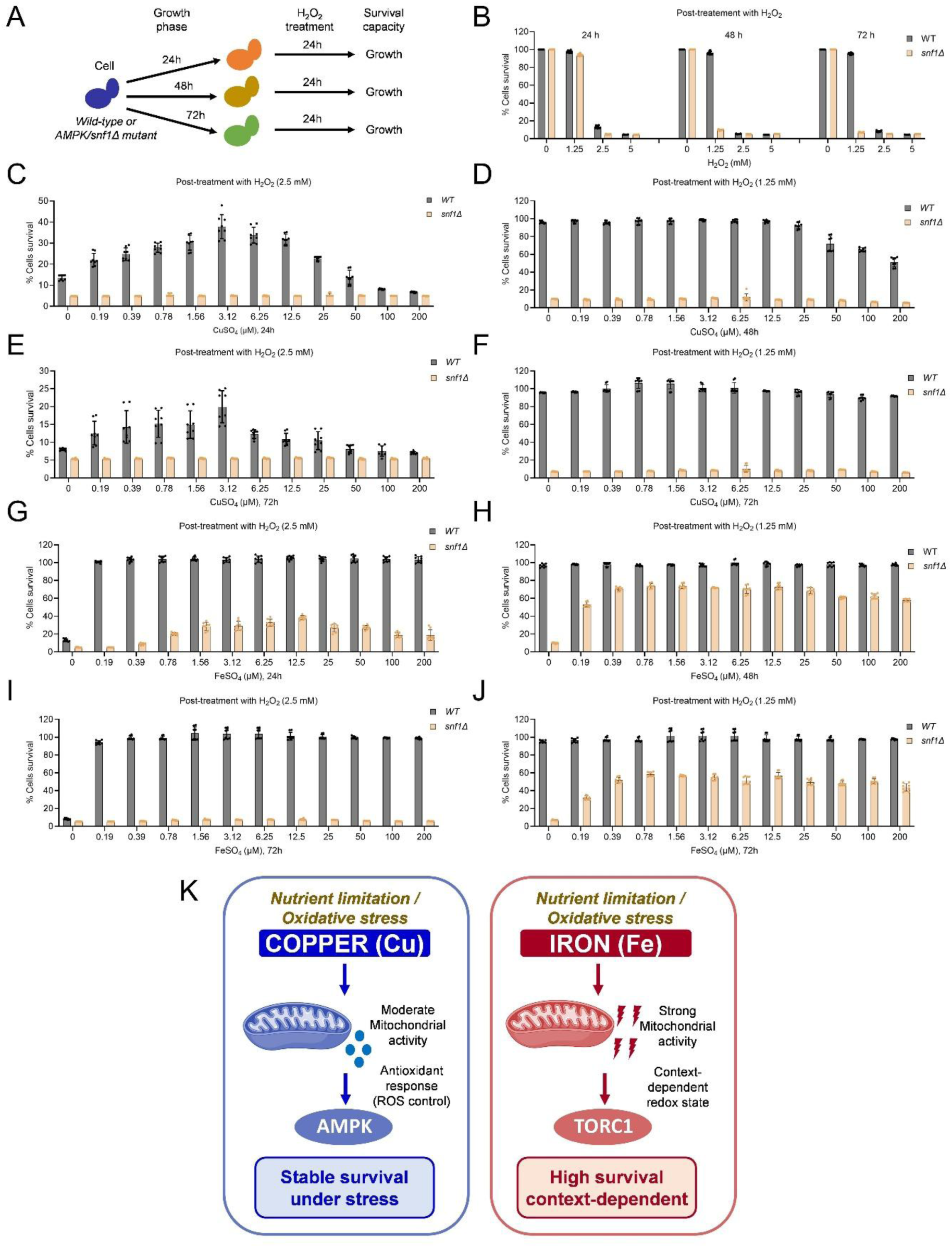
AMPK/Snf1 modulates copper- and iron-dependent oxidative stress tolerance. (A) Schematic representation of the experimental design for oxidative stress assays. Wild-type (WT) and *snf1Δ* cells were cultured for 24, 48, or 72 h, followed by H_2_O_2_ treatment and subsequent assessment of survival by outgrowth. (B) Survival of WT and *snf1Δ* cells following exposure to increasing concentrations of H_2_O_2_ (0–5 mM) after 24, 48, or 72 h of growth. Viability was assessed using a liquid outgrowth assay and expressed relative to untreated controls. Data represent mean ± SD (n = 16). (C–F) Survival of WT and *snf1Δ* cells pre-cultured with increasing concentrations of CuSO_4_ (0–200 µM) for 24 h (C), 48 h (D), or 72 h (E–F), followed by treatment with H_2_O_2_ (2.5 mM in C and E; 1.25 mM in D and F). Viability was assessed using a liquid outgrowth assay. Data represent mean ± SD (n = 18). (G–J) Survival of WT and *snf1Δ* cells pre-cultured with increasing concentrations of FeSO_4_ (0–200 µM) for 24 h (G), 48 h (H), or 72 h (I–J), followed by treatment with H_2_O_2_ (2.5 mM in G and I; 1.25 mM in H and J). Viability was assessed using a liquid outgrowth assay. Data represent mean ± SD (n = 8). (K) Proposed model for copper- and iron-dependent survival regulation under nutrient limitation and oxidative stress. Copper promotes AMPK/Snf1-associated antioxidant adaptation and stable stress survival, whereas iron enhances mitochondrial activity and TORC1-associated context-dependent survival through distinct redox regulatory mechanisms.

We next examined whether pre-exposure to copper or iron modulates oxidative stress tolerance. In wild-type cells, low concentrations of copper conferred protection against H_2_O_2_-induced cytotoxicity, whereas higher concentrations became detrimental, consistent with a dose-dependent hormetic response **(Figures 7C–7F)**. This protective effect was attenuated in *snf1Δ* cells, indicating that copper-mediated oxidative stress resistance is at least partially dependent on Snf1 signaling.

In contrast, iron supplementation provided robust protection against H_2_O_2_-induced toxicity across the tested concentration range **(Figures 7G–7J)**. However, the dependence on Snf1 varied with stress severity. Under mild oxidative stress (1.25 mM H₂O₂), iron improved survival in both wild-type and *snf1Δ* cells, indicating a Snf1-independent protective effect. In contrast, under more severe stress (2.5 mM H₂O₂), iron-mediated protection was markedly reduced in *snf1Δ* cells, demonstrating that Snf1 becomes essential under high-stress conditions.

These results reveal distinct modes of redox adaptation: copper confers a dose-dependent, Snf1-associated protective effect, whereas iron provides broader protection that becomes increasingly dependent on Snf1 as oxidative stress intensifies. These results position Snf1/AMPK as a critical regulator integrating metal-dependent stress responses to support cellular survival.

Collectively, these findings define distinct metal-dependent survival strategies under nutrient and stress conditions. Iron-mediated survival is closely linked to mitochondrial function and sustained TORC1 activity, whereas copper promotes survival through a more flexible, partially TORC1-independent mechanism that engages Snf1/AMPK signaling and antioxidant defense **(Figure 7K)**. These results establish a framework in which micronutrient availability shapes cellular state transitions between growth and survival through coordinated metabolic and signaling networks.

## DISCUSSION

Copper and iron are essential redox-active micronutrients that support cellular metabolism and stress adaptation ^3–13^. While their individual roles in redox biology are well established, how they interact to regulate cellular survival across distinct metabolic states remains unclear. Here, using nutrient-limited conditions that impose metabolic and redox stress, we show that copper and iron promote survival through convergent but mechanistically distinct pathways that are differentially constrained by nutrient signaling and stress-response networks.

A central finding of this study is that copper-mediated survival requires iron availability, indicating that copper does not act independently but instead modulates iron-dependent metabolic and redox processes. This is consistent with the well-established interdependence of copper and iron homeostasis, including copper-dependent iron uptake and utilization pathways ^3,6,9,19,20,71^. These findings suggest that cellular survival under stress depends on coordinated regulation of these micronutrients rather than their individual abundance.

Despite this coupling, copper and iron engage fundamentally different survival programs. Iron-dependent survival is tightly linked to TORC1 signaling and is diminished when TORC1–Sch9 activity is disrupted, indicating that iron requires a TORC1-permissive metabolic state. Consistent with this, iron sustains TORC1 activity during the transition to stationary phase. In contrast, copper exerts minimal effects on TORC1 activity and retains pro-survival capacity under TORC1 inhibition, indicating a more flexible mechanism that operates independently of canonical nutrient signaling constraints.

This distinction is further highlighted by their interaction with rapamycin and calorie restriction. Iron-mediated survival is attenuated under TORC1 inhibition, whereas copper remains compatible and even cooperative under strong TORC1 suppression. These findings indicate that survival can be achieved through multiple metabolic configurations: an iron-dependent state linked to sustained TORC1 activity and a copper-dependent state that aligns with TORC1-restricted, stress-adaptive conditions.

In addition to nutrient signaling, we identify AMPK/Snf1 as a key regulator of metal-dependent stress adaptation. Copper-mediated protection against oxidative stress is largely Snf1-dependent, whereas iron-mediated protection exhibits a stress-threshold dependence on Snf1 activity. Under mild stress, iron acts independently of Snf1, but under severe oxidative stress, Snf1 becomes essential. These findings position AMPK/Snf1 as a critical integrator of metal-dependent redox responses, enabling adaptation when stress exceeds a defined threshold^45,49–51,62,108^.

Our findings further indicate that metal-dependent survival extends beyond classical nutrient signaling to include resilience against organelle dysfunction. Disruption of vacuolar function using concanamycin, which has been shown to impair mitochondrial function and cellular lifespan, reduced survival, while both copper and iron restored viability under these conditions ^69,103–105^. This suggests that copper and iron can buffer the downstream consequences of organelle dysfunction, linking metal homeostasis to broader cellular stress resilience mechanisms.

Together, these results support a model in which cellular survival is governed by state-dependent integration of nutrient signaling and stress-response pathways. Within this framework, copper and iron differentially engage these regulatory axes: copper exhibits a dose-dependent, partially Snf1-dependent response that is compatible with reduced TORC1 signaling, whereas iron provides robust but context-dependent protection that requires TORC1 activity and, under high stress, Snf1 engagement. Thus, metal-dependent survival is not fixed but dynamically shaped by metabolic state and stress intensity.

The nutrient-limited conditions used in this study impose metabolic and redox constraints that capture key features of aging-associated stress, including reduced nutrient availability, altered redox balance, and increased reliance on adaptive signaling pathways. In aging systems, decline in AMPK signaling activity and dysregulation of iron homeostasis are well documented ^45,49–51,62,69,103–105,108,109^. Our findings suggest that such changes may shift the balance between beneficial and detrimental effects of iron, whereby insufficient iron limits metabolic capacity, while excess iron promotes oxidative damage, particularly when stress-response pathways are compromised.

These observations may also have implications for cancer biology. Iron metabolism is increasingly targeted therapeutically, particularly through ferroptosis-based strategies ^110–112^. However, our results indicate that iron can also support cellular survival under specific metabolic and signaling conditions. In nutrient-limited tumor microenvironments, where survival depends on metabolic flexibility and stress tolerance ^37–42^, iron availability may contribute to persistence rather than cell death. In this context, copper–iron interactions may further modulate cellular outcomes by influencing redox balance and metabolic adaptation.

Finally, our findings raise the possibility that sublethal or persistent stress conditions may promote long-term survival states that depend on metal availability and adaptive signaling. Cells that evade cell death under such conditions may adopt altered functional states, including senescence-like phenotypes, which have been associated with iron accumulation and redox imbalance in multiple systems ^113,114^. If such states are reversible, they may contribute to disease progression or relapse through re-entry into proliferative programs ^115–120^. Although these possibilities remain speculative, they highlight the importance of considering context-dependent effects of metal homeostasis in disease settings.

Whether these mechanisms are conserved in mammalian systems remains to be determined and warrants further investigation. Nevertheless, the established connections among iron metabolism, mitochondrial function, mTORC1 signaling, and stress adaptation suggest potential translational relevance ^6,7,9,30,37,40,41,57,76,77,116,121–127^. Future studies in human cells will be required to determine how copper–iron interactions influence cellular survival under defined metabolic and stress conditions.

In conclusion, this study defines copper and iron as context-dependent regulators of cellular survival that act through distinct but interconnected nutrient and stress signaling pathways. By integrating metal homeostasis with TORC1 and AMPK/Snf1 signaling, our work provides a framework for understanding how redox-active micronutrients shape cellular adaptation under stress, with potential implications for aging, metabolic disorders, and cancer.

## METHODS

### Yeast strains and growth conditions

The budding yeast *Saccharomyces cerevisiae* strains used in this study included the prototrophic CEN.PK113-7D ^128^ background and the auxotrophic BY4743 strain (obtained from Euroscarf). Gene knockout strains were constructed using a PCR-mediated homologous recombination strategy as previously described ^129^. Cells were recovered from glycerol stocks and streaked onto YPD agar plates composed of 1% yeast extract, 2% peptone, 2% glucose, and 2.5% agar, followed by incubation at 30 °C for 48–72 hours. For experimental assays, CEN.PK113-7D strains were propagated in synthetic defined (SD) medium containing 6.7 g/L yeast nitrogen base with ammonium sulfate (lacking amino acids) and supplemented with 2% glucose. For BY4743 auxotrophic strains, the same SD medium was additionally supplemented with histidine (40 mg/L), leucine (160 mg/L), and uracil (40 mg/L) to support growth.

### Chemical treatments

Cells were exposed to rapamycin, antimycin A and concanamycin A, which were prepared as concentrated stock solutions in dimethyl sulfoxide (DMSO). Control samples received an equivalent volume of DMSO to account for vehicle effects. The final concentration of DMSO in all yeast experiments was maintained at ≤1%. Metal salts, including CuSO_4_, CuCl_3_, FeSO_4_·7H_2_O, and FeCl_3_, as well as bathophenanthroline disulfonate (BPS), were dissolved in sterile water to generate stock solutions prior to use.

### Growth assay

Yeast growth in response to chemical treatments was quantified using a 96-well microplate–based format. Prototrophic CEN.PK113-7D and auxotrophic BY4743 *Saccharomyces cerevisiae* strains were maintained in synthetic defined (SD) medium consisting of 6.7 g/L yeast nitrogen base with ammonium sulfate (lacking amino acids) and supplemented with 2% glucose. For BY4743 strains, histidine (40 mg/L), leucine (160 mg/L), and uracil (40 mg/L) were added to the medium to support auxotrophic growth requirements. Cells were diluted to a starting density of approximately OD600 ∼0.2 and aliquoted into 96-well plates at 200 µL per well. Test compounds were applied as serial two-fold dilutions across the plate. Cultures were incubated at 30°C, and cell density was determined by measuring absorbance at 600nm (OD600) using a microplate reader at designated time points.

### Chronological aging survival assay

Cellular survival was evaluated using the chronological lifespan (CLS) assay as described previously ^63,65,74^. *Saccharomyces cerevisiae* prototrophic CEN.PK113-7D strains were cultured in synthetic defined (SD) medium containing 6.7 g/L yeast nitrogen base with ammonium sulfate (without amino acids) supplemented with 2% glucose. SD medium supplemented with histidine (40 mg/L), leucine (160 mg/L), and uracil (40 mg/L) for auxotrophic BY4743 strains. To initiate CLS experiments, overnight cultures grown at 30 °C with shaking at 220 rpm were diluted to an initial optical density at 600 nm (OD600) of approximately 0.2 in fresh SD medium. CLS was assessed using two complementary approaches. (i) Microplate outgrowth assay: Cells were aged in 96-well plates containing 200 µL SD medium per well at 30 °C. At indicated time points, 2 µL of stationary-phase culture was transferred into a second 96-well plate containing 200 µL YPD medium and incubated for 24 h at 30 °C. Outgrowth was quantified by measuring OD600 using a microplate reader. (ii) Spot dilution viability assay: Cells were aged in glass flasks containing SD medium at 30 °C with shaking at 220 rpm. At indicated time points, cultures were collected, washed, and normalized to OD600 = 1.0 in YPD medium. Cells were serially diluted 10-fold in YPD, and 3 µL of each dilution was spotted onto YPD agar plates. Plates were incubated for 48 h at 30 °C, and colony outgrowth was documented using a GelDoc imaging system.

### Chronological lifespan assay

Cell survival during chronological aging was assessed using established CLS methodologies ^63,65,74^. Prototrophic CEN.PK113-7D and auxotrophic BY4743 *Saccharomyces cerevisiae* strains were cultured in synthetic defined (SD) medium composed of 6.7 g/L yeast nitrogen base with ammonium sulfate (without amino acids) and supplemented with 2% glucose. For BY4743 strains, the medium was additionally supplemented with histidine (40 mg/L), leucine (160 mg/L), and uracil (40 mg/L). For CLS experiments, overnight cultures grown at 30°C with shaking (220 rpm) were diluted to an initial density of approximately OD600 ∼0.2 in fresh SD medium and maintained under aging conditions. Survival was evaluated using two complementary approaches:

i. Microplate-based outgrowth assay: Cells were aged in 96-well plates containing 200 µL SD medium per well at 30 °C. At specified time points, 2 µL of aged culture was transferred into fresh 96-well plates containing 200 µL YPD medium and incubated for 24 h at 30 °C. Regrowth capacity, as a proxy for viability, was determined by measuring OD600 using a microplate reader.
ii. Spot dilution viability assay: In parallel, cultures were aged in SD medium in shaking flasks at 30 °C (220 rpm). At indicated time points, cells were harvested, washed, and adjusted to OD600 = 1.0 in YPD medium. Serial 10-fold dilutions were prepared, and 3 µL of each dilution was spotted onto YPD agar plates. Plates were incubated at 30 °C for 48 h, and colony formation was recorded using a GelDoc imaging system.

### Combined rapamycin, calorie restriction, and metal supplementation assay

The combined effects of rapamycin (RM), calorie restriction (CR), and metal supplementation (copper or iron) on yeast growth and survival were assessed using modified chronological lifespan (CLS) assays, as described previously ^66^. Prototrophic CEN.PK113-7D Saccharomyces cerevisiae cells were cultured in synthetic defined (SD) medium containing 6.7 g/L yeast nitrogen base with ammonium sulfate (without amino acids) supplemented with 2% glucose. Calorie restriction conditions were established by reducing glucose concentration to 0.5%. Cells were inoculated at an initial density of approximately OD600 ∼0.2 and grown under the indicated treatment conditions, including rapamycin and either copper or iron supplementation. Growth kinetics were monitored at defined time points prior to entry into late stationary phase. Cell survival during chronological aging was subsequently assessed using outgrowth-based assays at indicated time points. Two complementary methods were employed:

i. Microplate outgrowth assay: Cells were aged in 96-well plates containing 200 µL SD medium per well at 30 °C. At specified time points, 2 µL of aged cultures was transferred into fresh 96-well plates containing 200 µL YPD medium and incubated for 24 h at 30 °C. Regrowth was quantified by measuring OD600 using a microplate reader.
ii. Outgrowth dilution assay: In parallel, cells were cultured and aged in flasks containing SD medium at 30 °C with shaking (220 rpm). At indicated chronological time points, cultures were harvested, washed, and normalized to OD600 = 1.0 in YPD medium. Cells were then serially diluted 10-fold in YPD in 96-well plates and incubated for 24 h at 30 °C. Outgrowth was determined by measuring OD600 using a microplate reader.

### Oxidative stress growth assay

To evaluate the impact of oxidative stress on copper- and iron-treated cells, hydrogen peroxide (H_2_O_2_) challenge assays were performed. Prototrophic CEN.PK113-7D *Saccharomyces cerevisiae* cells were cultured in synthetic defined (SD) medium containing 6.7 g/L yeast nitrogen base with ammonium sulfate (without amino acids) supplemented with 2% glucose. Cells were inoculated at an initial density of approximately OD600 ∼0.2 and cultured in the presence or absence of copper or iron for 24, 48, and 72 hours. At each time point, aliquots of the pre-incubated cultures were harvested and exposed to increasing concentrations of H_2_O_2_ to induce oxidative stress. Following H_2_O_2_ treatment, cells were incubated for an additional 24 hours, and viability was assessed using a 96-well microplate–based growth assay. Cell density was quantified by measuring OD600 using a microplate reader, providing a readout of survival following oxidative stress exposure.

### RNA extraction and quantitative real-time PCR

Total RNA was isolated using the RNeasy Mini Kit (Qiagen) following the manufacturer’s instructions and as described previously ^64,66,130^. Mechanical disruption was included to ensure efficient cell lysis. RNA concentration and purity were assessed using a NanoDrop 2000 spectrophotometer. cDNA synthesis was performed using 1 μg of total RNA with the QuantiTect Reverse Transcription Kit (Qiagen). Control reactions lacking either reverse transcriptase or RNA template were included to confirm the absence of genomic DNA contamination and reagent background. Quantitative real-time PCR (qRT-PCR) was carried out in a final reaction volume of 20 μL containing 20 ng of cDNA using the SYBR Fast Universal qPCR Kit (Kapa Biosystems). Amplification was performed on a QuantStudio 6 Flex system (Applied Biosystems) under the following cycling conditions: 95 °C for 180 s, followed by 40 cycles of 95 °C for 1 s and 60 °C for 20 s. Melt curve analysis was conducted after amplification to verify product specificity and exclude primer-dimer formation. Gene expression levels were normalized to the housekeeping gene *ACT1*, and relative expression changes were calculated using the 2^−ΔΔCt^ method as previously described ^64,130^.

### Mitochondrial membrane potential measurement

Mitochondrial membrane potential was assessed in yeast cells using the fluorescent dye DiOC_6_, as described previously ^67,131^. Cells were collected, washed with phosphate-buffered saline (PBS), and incubated with 50 nM DiOC_6_ at 30 °C for 30 min in the dark. Following staining, cells were washed to remove excess dye and resuspended in PBS for fluorescence measurement. Fluorescence was recorded using excitation and emission wavelengths of 482 nm and 504 nm, respectively. Signal intensity was normalized to cell density (OD600) to account for differences in cell number.

### TORC1 activity assay

TORC1 activity was evaluated by monitoring phosphorylation of the downstream target Sch9, as described previously with minor modifications ^130,132^. Prototrophic *Saccharomyces cerevisiae* CEN.PK113-7D wild-type cells expressing Sch9-6×HA were cultured in SD medium and subjected to the indicated treatment conditions. Cells were harvested at specified time points for protein extraction. Protein samples were separated by SDS–PAGE and transferred onto nitrocellulose membranes for immunoblot analysis. Membranes were blocked with 5% (w/v) non-fat milk in TBS containing 0.1% Tween-20 (TBST). Phosphorylated Sch9 was detected using an anti-HA 3F10 antibody (1:2000; Roche Life Science), followed by incubation with an HRP-conjugated goat anti-rat secondary antibody (1:5000; Santa Cruz Biotechnology). Signal detection was performed using ECL Prime reagent (Amersham), and images were acquired using an iBright CL1500 imaging system (Thermo Fisher Scientific).

### Statistical analysis

All statistical analyses and data visualization were performed using GraphPad Prism (version 11). Data are presented as mean ± standard deviation (SD) unless otherwise stated. Statistical significance was determined using Student’s t-test, one-way ANOVA, or two-way ANOVA, as appropriate, with multiple comparison corrections applied where indicated in the figure legends. Significance levels were defined as *P < 0.05, **P < 0.01, ***P < 0.001, and ****P < 0.0001, while “ns” denotes not statistically significant.

## Code availability

No custom code or mathematical algorithm was used in this study.

## Data availability

Further information and requests for resources and reagents should be directed to and will be fulfilled by the Lead Contact, Dr. Mohammad Alfatah.

## ACKNOWLEDGMENTS

This work is supported by the Young Investigator Research Grant (YIRG), National Medical Research Council, Singapore (MOH-001348-00) and US NAM Healthy Longevity Catalyst Awards Grant (MOH-001439). Arshia Naaz is supported by A*STAR-CDF grant (C243512027).

## AUTHOR CONTRIBUTIONS STATEMENT

Arshia Naaz: Writing-original draft, Investigation, Formal analysis, Funding Acquisition

Trishia Cheng Yi Ning: Investigation, Formal analysis

Mingtong Gao: Investigation, Formal analysis

Jovian Jing Lin: Investigation, Formal analysis Rajkumar Dorajoo: Review and editing

Brian K Kennedy: Review and editing

Mohammad Alfatah: Conceptualization, Writing-review and editing, Funding acquisition.

All authors read, critically reviewed and approved the final manuscript.

M.A is the guarantor of this work.

## REFERENCES

1. Zheng, L. et al. Dietary Copper Intake and Biological Aging Among US Adults, NHANES 2003–2018. Aging Cell 24, e70272 (2025).

2. Yang, L., Chen, X., Cheng, H. & Zhang, L. Dietary Copper Intake and Risk of Stroke in Adults: A Case-Control Study Based on National Health and Nutrition Examination Survey 2013–2018. Nutrients 14, (2022).

3. Kaplan, J. & Ward, D. M. The essential nature of iron usage and regulation. Current Biology vol. 23 R642–R646 at 10.1016/j.cub.2013.05.033 (2013).

4. Guthrie, L. M. et al. Elesclomol alleviates Menkes pathology and mortality by escorting Cu to cuproenzymes in mice. Science (80-.). 368, 620–625 (2020).

5. Kauppila, T. E. S., Kauppila, J. H. K. & Larsson, N. G. Mammalian Mitochondria and Aging: An Update. Cell Metabolism at 10.1016/j.cmet.2016.09.017 (2017).

6. Festa, R. A. & Thiele, D. J. Copper: An essential metal in biology. Current Biology vol. 21 R877–R883 at 10.1016/j.cub.2011.09.040 (2011).

7. Teh, M. R., Armitage, A. E. & Drakesmith, H. Why cells need iron: a compendium of iron utilisation. Trends in Endocrinology and Metabolism vol. 35 1026–1049 at 10.1016/j.tem.2024.04.015 (2024).

8. Lane, A. R., Roberts, B. R., Fahrni, C. J. & Faundez, V. A primer on copper biology in the brain. Neurobiology of Disease vol. 212 106974 at 10.1016/j.nbd.2025.106974 (2025).

9. Xu, W., Barrientos, T. & Andrews, N. C. Iron and copper in mitochondrial diseases. Cell Metabolism vol. 17 319–328 at 10.1016/j.cmet.2013.02.004 (2013).

10. Ruiz, L. M., Libedinsky, A. & Elorza, A. A. Role of Copper on Mitochondrial Function and Metabolism. Frontiers in Molecular Biosciences vol. 8 711227 at 10.3389/fmolb.2021.711227 (2021).

11. MUNOMETa, I. M., et al. Copper supports regulatory T cell energetic state to sustain peripheral immune tolerance. Sci. Immunol. 11, 2573 (2026).

12. Chen, L. et al. Homeostasis and metabolism of iron and other metal ions in neurodegenerative diseases. Signal Transduction and Targeted Therapy vol. 10 31-at 10.1038/s41392-024-02071-0 (2025).

13. Rouault, T. A. & Tong, W. H. Iron-sulphur cluster biogenesis and mitochondrial iron homeostasis. Nature Reviews Molecular Cell Biology vol. 6 345–351 at 10.1038/nrm1620 (2005).

14. Sun, N., Youle, R. J. & Finkel, T. The Mitochondrial Basis of Aging. Molecular Cell at 10.1016/j.molcel.2016.01.028 (2016).

15. Liguori, I., et al. Oxidative stress, aging, and diseases. Clinical Interventions in Aging at 10.2147/CIA.S158513 (2018).

16. Yu, P.-L. & Xu, C.-R. Vitamin C slows primate aging by targeting iron-driven lipid peroxidation. Cell Metab. 38, 635–637 (2026).

17. Chandimali, N. et al. Free radicals and their impact on health and antioxidant defenses: a review. Cell Death Discovery vol. 11 19- at 10.1038/s41420-024-02278-8 (2025).

18. Brewer, G. J. Risks of copper and iron toxicity during aging in humans. Chemical Research in Toxicology vol. 23 319–326 at 10.1021/tx900338d (2010).

19. Collins, J. F., Prohaska, J. R. & Knutson, M. D. Metabolic crossroads of iron and copper. Nutrition Reviews vol. 68 133–147 at 10.1111/j.1753-4887.2010.00271.x (2010).

20. Taylor, A. B., Stoj, C. S., Ziegler, L., Kosman, D. J. & Hart, P. J. The copper-iron connection in biology: Structure of the metallo-oxidase Fet3p. Proc. Natl. Acad. Sci. U. S. A. 102, 15459–15464 (2005).

21. Gulec, S. & Collins, J. F. Molecular mediators governing iron-copper interactions. Annu. Rev. Nutr. 34, 95–116 (2014).

22. Lewis, M. S. Iron and copper in the treatment of anemia in children. J. Am. Med. Assoc. 96, 1135–1138 (1931).

23. Doguer, C., Ha, J. H. & Collins, J. F. Intersection of Iron and Copper Metabolism in the Mammalian Intestine and Liver. Compr. Physiol. 8, 1433–1461 (2018).

24. Galy, B. et al. Iron regulatory proteins secure mitochondrial iron sufficiency and function. Cell Metab. (2010) doi:10.1016/j.cmet.2010.06.007.

25. Dutt, S., Hamza, I. & Bartnikas, T. B. Molecular Mechanisms of Iron and Heme Metabolism. Annual Review of Nutrition vol. 42 311–335 at 10.1146/annurev-nutr-062320-112625 (2022).

26. Martínez-Reyes, I. & Chandel, N. S. Mitochondrial TCA cycle metabolites control physiology and disease. Nature Communications vol. 11 at 10.1038/s41467-019-13668-3 (2020).

27. Lin, S. J., Pufahl, R. A., Dancis, A., O’Halloran, T. V. & Culotta, V. C. A role for the Saccharomyces cerevisiae ATX1 gene in copper trafficking and iron transport. J. Biol. Chem. (1997) doi:10.1074/jbc.272.14.9215.

28. Neidlein, S., Wirth, R. & Pourhassan, M. Iron deficiency, fatigue and muscle strength and function in older hospitalized patients. Eur. J. Clin. Nutr. (2021) doi:10.1038/s41430-020-00742-z.

29. Chen, L., Min, J. & Wang, F. Copper homeostasis and cuproptosis in health and disease. Signal Transduction and Targeted Therapy vol. 7 378- at 10.1038/s41392-022-01229-y (2022).

30. Müller, S., Cañeque, T., Solier, S. & Rodriguez, R. Copper and iron orchestrate cell-state transitions in cancer and immunity. Trends in Cell Biology vol. 35 105–114 at 10.1016/j.tcb.2024.07.005 (2025).

31. Pasricha, S. R., Tye-Din, J., Muckenthaler, M. U. & Swinkels, D. W. Iron deficiency. The Lancet at 10.1016/S0140-6736(20)32594-0 (2021).

32. Cappellini, M. D., Musallam, K. M. & Taher, A. T. Iron deficiency anaemia revisited. Journal of Internal Medicine at 10.1111/joim.13004 (2020).

33. Beard, J. L. Iron biology in immune function, muscle metabolism and neuronal functioning. in Journal of Nutrition (2001). doi:10.1093/jn/131.2.568s.

34. Busti, F., Campostrini, N., Martinelli, N. & Girelli, D. Iron deficiency in the elderly population, revisited in the hepcidin era. Frontiers in Pharmacology at 10.3389/fphar.2014.00083 (2014).

35. López-Otín, C., Blasco, M. A., Partridge, L., Serrano, M. & Kroemer, G. Hallmarks of aging: An expanding universe. Cell 186, 243–278 (2023).

36. López-Otín, C., Blasco, M. A., Partridge, L., Serrano, M. & Kroemer, G. The hallmarks of aging. Cell 153, 1194–217 (2013).

37. Abbott, K. L. et al. Nutrient requirements of organ-specific metastasis in breast cancer. Nature 649, 1292–1301 (2026).

38. Lobel, G. P., Jiang, Y. & Simon, M. C. Tumor microenvironmental nutrients, cellular responses, and cancer. Cell Chemical Biology vol. 30 1015–1032 at 10.1016/j.chembiol.2023.08.011 (2023).

39. Vaziri-Gohar, A. et al. Limited nutrient availability in the tumor microenvironment renders pancreatic tumors sensitive to allosteric IDH1 inhibitors. Nat. Cancer 3, 852–865 (2022).

40. Hanahan, D. Hallmarks of cancer-Then and now, and beyond. Cell 0, (2026).

41. Finicle, B. T., Jayashankar, V. & Edinger, A. L. Nutrient scavenging in cancer. Nature Reviews Cancer vol. 18 619–633 at 10.1038/s41568-018-0048-x (2018).

42. Cognet, G. & Muir, A. Identifying metabolic limitations in the tumor microenvironment. Science Advances vol. 10 eadq7305 at 10.1126/sciadv.adq7305 (2024).

43. Goul, C., Peruzzo, R. & Zoncu, R. The molecular basis of nutrient sensing and signalling by mTORC1 in metabolism regulation and disease. Nature Reviews Molecular Cell Biology vol. 24 857–875 at 10.1038/s41580-023-00641-8 (2023).

44. Kim, J. & Guan, K. L. mTOR as a central hub of nutrient signalling and cell growth. Nature Cell Biology vol. 21 63–71 at 10.1038/s41556-018-0205-1 (2019).

45. Hardie, D. G., Ross, F. A. & Hawley, S. A. AMPK: A nutrient and energy sensor that maintains energy homeostasis. Nature Reviews Molecular Cell Biology at 10.1038/nrm3311 (2012).

46. González, A., Hall, M. N., Lin, S. C. & Hardie, D. G. AMPK and TOR: The Yin and Yang of Cellular Nutrient Sensing and Growth Control. Cell Metabolism at 10.1016/j.cmet.2020.01.015 (2020).

47. Zhang, C. S. et al. Fructose-1,6-bisphosphate and aldolase mediate glucose sensing by AMPK. Nature (2017) doi:10.1038/nature23275.

48. Weir, H. J. et al. Dietary Restriction and AMPK Increase Lifespan via Mitochondrial Network and Peroxisome Remodeling. Cell Metab. (2017) doi:10.1016/j.cmet.2017.09.024.

49. Hedbacker, K. & Carlson, M. SNF1/AMPK pathways in yeast. Frontiers in Bioscience at 10.2741/2854 (2008).

50. Garcia, D. & Shaw, R. J. AMPK: Mechanisms of Cellular Energy Sensing and Restoration of Metabolic Balance. Molecular Cell at 10.1016/j.molcel.2017.05.032 (2017).

51. Burkewitz, K., Zhang, Y. & Mair, W. B. AMPK at the nexus of energetics and aging. Cell Metabolism at 10.1016/j.cmet.2014.03.002 (2014).

52. Lin, S. C. & Hardie, D. G. AMPK: Sensing Glucose as well as Cellular Energy Status. Cell Metabolism at 10.1016/j.cmet.2017.10.009 (2018).

53. Hallett, J. E. H. et al. Snf1/AMPK promotes the formation of Kog1/Raptor-bodies to increase the activation threshold of TORC1 in budding yeast. Elife 4, e09181 (2015).

54. Schmauck-Medina, T. et al. Dietary restriction in aging and longevity. Nat. Aging 2026 1–21 (2026) doi:10.1038/s43587-026-01091-5.

55. Saxton, R. A. & Sabatini, D. M. mTOR signaling in growth, metabolism, and disease. Cell 168, 960–976 (2017).

56. Sabatini, D. M. Twenty-five years of mTOR: Uncovering the link from nutrients to growth. Proc. Natl. Acad. Sci. U. S. A. 114, 11818–11825 (2017).

57. Liu, G. Y. & Sabatini, D. M. mTOR at the nexus of nutrition, growth, ageing and disease. Nat. Rev. Mol. Cell Biol. 21, 1–21 (2020).

58. Kennedy, B. K. & Lamming, D. W. The Mechanistic Target of Rapamycin: The Grand ConducTOR of Metabolism and Aging. Cell Metabolism at 10.1016/j.cmet.2016.05.009 (2016).

59. Loewith, R. & Hall, M. N. Target of rapamycin (TOR) in nutrient signaling and growth control. Genetics 189, 1177–1201 (2011).

60. González, A. & Hall, M. N. Nutrient sensing and TOR signaling in yeast and mammals. EMBO J. 36, 397–408 (2017).

61. Dibble, C. C. & Manning, B. D. Signal integration by mTORC1 coordinates nutrient input with biosynthetic output. Nat. Cell Biol. 15, 555–564 (2013).

62. Herzig, S. & Shaw, R. J. AMPK: Guardian of metabolism and mitochondrial homeostasis. Nature Reviews Molecular Cell Biology at 10.1038/nrm.2017.95 (2018).

63. Longo, V. D., Shadel, G. S., Kaeberlein, M. & Kennedy, B. Replicative and chronological aging in saccharomyces cerevisiae. Cell Metabolism at 10.1016/j.cmet.2012.06.002 (2012).

64. Alfatah, M. et al. Uncharacterized yeast gene YBR238C, an effector of TORC1 signaling in a mitochondrial feedback loop, accelerates cellular aging via HAP4- and RMD9-dependent mechanisms. eLife 12:RP92178 10.7554/eLife.92178.1 (2023).

65. Alfatah, M. & Eisenhaber, F. The PICLS high-throughput screening method for agents extending cellular longevity identifies 2,5-anhydro-D-mannitol as novel anti-aging compound. GeroScience 45, 141–158 (2022).

66. Zhang, Y. et al. Systematic transcriptomics analysis of calorie restriction and rapamycin unveils their synergistic interaction in prolonging cellular lifespan. Commun. Biol. 8, 1–16 (2025).

67. Jing, J. L., Ning, T. C. Y., Natali, F., Eisenhaber, F. & Alfatah, M. Iron Supplementation Delays Aging and Extends Cellular Lifespan through Potentiation of Mitochondrial Function. Cells 11, 862 (2022).

68. Shen, H. An IRON-clad Connection between Aging Organelles. Cell at 10.1016/j.cell.2019.12.037 (2020).

69. Hughes, C. E. et al. Cysteine Toxicity Drives Age-Related Mitochondrial Decline by Altering Iron Homeostasis. Cell (2020) doi:10.1016/j.cell.2019.12.035.

70. Cheng, C. et al. Histone H3 cysteine 110 enhances iron metabolism and modulates replicative life span in Saccharomyces cerevisiae. Sci. Adv. (2025) doi:10.1126/sciadv.adv4082.

71. Hassett, R. F., Romeo, A. M. & Kosman, D. J. Regulation of high affinity iron uptake in the yeast Saccharomyces cerevisiae: Role of dioxygen and Fe(II). J. Biol. Chem. 273, 7628–7636 (1998).

72. Nolfi-Donegan, D., Braganza, A. & Shiva, S. Mitochondrial electron transport chain: Oxidative phosphorylation, oxidant production, and methods of measurement. Redox Biology at 10.1016/j.redox.2020.101674 (2020).

73. Li, J., Kim, S. G. & Blenis, J. Rapamycin: One drug, many effects. Cell Metabolism vol. 19 373–379 at 10.1016/j.cmet.2014.01.001 (2014).

74. Powers, R. W., Kaeberlein, M., Caldwell, S. D., Kennedy, B. K. & Fields, S. Extension of chronological life span in yeast by decreased TOR pathway signaling. Genes Dev. 20, 174–184 (2006).

75. Shapiro, J. S. et al. Iron drives anabolic metabolism through active histone demethylation and mTORC1. Nat. Cell Biol. (2023) doi:10.1038/s41556-023-01225-6.

76. Carney, E. F. A new pathway links iron sensing with histone demethylation and regulation of mTORC1 activity. Nat. Rev. Nephrol. 19, 753 (2023).

77. Bayeva, M. et al. MTOR regulates cellular iron homeostasis through tristetraprolin. Cell Metab. 16, 645–657 (2012).

78. Knight, Z. A., Schmidt, S. F., Birsoy, K., Tan, K. & Friedman, J. M. A critical role for mTORC1 in erythropoiesis and anemia. Elife 3, e01913 (2014).

79. Conjard-Duplany, A. et al. Muscle mTOR controls iron homeostasis and ferritinophagy via NRF2, HIFs and AKT/PKB signaling pathways. Cell. Mol. Life Sci. 82, 178- (2025).

80. Watson, A., Lipina, C., McArdle, H. J., Taylor, P. M. & Hundal, H. S. Iron depletion suppresses mTORC1-directed signalling in intestinal Caco-cells via induction of REDD1. Cell. Signal. 28, 412–424 (2016).

81. Huang, H. et al. Two mTOR inhibitors, rapamycin and Torin 1, differentially regulate iron-induced generation of mitochondrial ROS. BioMetals 30, 975–980 (2017).

82. Zhou, X. et al. Cytochrome b561 regulates iron metabolism by activating the Akt/mTOR pathway to promote Breast Cancer Cells proliferation. Exp. Cell Res. 431, 113760 (2023).

83. Bogdan, A. R., Miyazawa, M., Hashimoto, K. & Tsuji, Y. Regulators of Iron Homeostasis: New Players in Metabolism, Cell Death, and Disease. Trends in Biochemical Sciences vol. 41 274–286 at 10.1016/j.tibs.2015.11.012 (2016).

84. Fontana, L. & Partridge, L. Promoting health and longevity through diet: From model organisms to humans. Cell at 10.1016/j.cell.2015.02.020 (2015).

85. Colman, R. J. et al. Caloric restriction delays disease onset and mortality in rhesus monkeys. Science (80-.). 325, 201–204 (2009).

86. Turturro, A. et al. Growth curves and survival characteristics of the animals used in the biomarkers of aging program. Journals Gerontol. - Ser. A Biol. Sci. Med. Sci. 54, B492–B501 (1999).

87. Acosta-Rodríguez, V. et al. Circadian alignment of early onset caloric restriction promotes longevity in male C57BL/6J mice. Science (80-.). 376, 1192–1202 (2022).

88. Mair, W. & Dillin, A. Aging and survival: The genetics of life span extension by dietary restriction. Annual Review of Biochemistry vol. 77 727–754 at 10.1146/annurev.biochem.77.061206.171059 (2008).

89. Igarashi, M. & Guarente, L. mTORC1 and SIRT1 Cooperate to Foster Expansion of Gut Adult Stem Cells during Calorie Restriction. Cell 166, 436–450 (2016).

90. Tulsian, R., Velingkaar, N. & Kondratov, R. Caloric restriction effects on liver mTOR signaling are time-of-day dependent. Aging (Albany. NY). 10, 1640–1648 (2018).

91. Ferreira-Marques, M., Carvalho, A., Cavadas, C. & Aveleira, C. A. PI3K/AKT/MTOR and ERK1/2-MAPK signaling pathways are involved in autophagy stimulation induced by caloric restriction or caloric restriction mimetics in cortical neurons. Aging (Albany. NY). 13, 7872–7882 (2021).

92. Nicastro, R., Sardu, A., Panchaud, N. & De Virgilio, C. The architecture of the Rag GTPase signaling network. Biomolecules at 10.3390/biom7030048 (2017).

93. Hatakeyama, R. & De Virgilio, C. Unsolved mysteries of Rag GTPase signaling in yeast. Small GTPases at 10.1080/21541248.2016.1211070 (2016).

94. Kira, S. et al. Dynamic relocation of the TORC1-Gtr1/2-Ego1/2/3 complex is regulated by Gtr1 and Gtr2. Mol. Biol. Cell 27, 382–396 (2016).

95. Binda, M. et al. The Vam6 GEF Controls TORC1 by Activating the EGO Complex. Mol. Cell (2009) doi:10.1016/j.molcel.2009.06.033.

96. Bonfils, G. et al. Leucyl-tRNA Synthetase Controls TORC1 via the EGO Complex. Mol. Cell (2012) doi:10.1016/j.molcel.2012.02.009.

97. Péli-Gulli, M. P., Sardu, A., Panchaud, N., Raucci, S. & De Virgilio, C. Amino Acids Stimulate TORC1 through Lst4-Lst7, a GTPase-Activating Protein Complex for the Rag Family GTPase Gtr2. Cell Rep. (2015) doi:10.1016/j.celrep.2015.08.059.

98. Urban, J. et al. Sch9 Is a Major Target of TORC1 in Saccharomyces cerevisiae. Mol. Cell 26, 663–674 (2007).

99. Fabrizio, P., Pozza, F., Pletcher, S. D., Gendron, C. M. & Longo, V. D. Regulation of longevity and stress resistance by Sch9 in yeast. Science (80-.). 292, 288–290 (2001).

100. Kapahi, P. et al. With TOR, less is more: A key role for the conserved nutrient-sensing TOR pathway in aging. Cell Metabolism vol. 11 453–465 at 10.1016/j.cmet.2010.05.001 (2010).

101. Wierman, M. B., Maqani, N., Strickler, E., Li, M. & Smith, J. S. Caloric Restriction Extends Yeast Chronological Life Span by Optimizing the Snf1 (AMPK) Signaling Pathway. Mol. Cell. Biol. (2017) doi:10.1128/mcb.00562-16.

102. Huang, L. S., Cobessi, D., Tung, E. Y. & Berry, E. A. Binding of the respiratory chain inhibitor antimycin to the mitochondrial bc1 complex: A new crystal structure reveals an altered intramolecular hydrogen-bonding pattern. J. Mol. Biol. 351, 573–597 (2005).

103. Cotter, K., Stransky, L., McGuire, C. & Forgac, M. Recent Insights into the Structure, Regulation, and Function of the V-ATPases. Trends in Biochemical Sciences vol. 40 611–622 at 10.1016/j.tibs.2015.08.005 (2015).

104. Hashmi, F. & Kane, P. M. V-ATPase Disassembly at the Yeast Lysosome-Like Vacuole Is a Phenotypic Driver of Lysosome Dysfunction in Replicative Aging. Aging Cell 24, e14487 (2025).

105. Colacurcio, D. J. & Nixon, R. A. Disorders of lysosomal acidification—The emerging role of v-ATPase in aging and neurodegenerative disease. Ageing Research Reviews vol. 32 75–88 at 10.1016/j.arr.2016.05.004 (2016).

106. Pan, Y., Schroeder, E. A., Ocampo, A., Barrientos, A. & Shadel, G. S. Regulation of yeast chronological life span by TORC1 via adaptive mitochondrial ROS signaling. Cell Metab. (2011) doi:10.1016/j.cmet.2011.03.018.

107. Bonawitz, N. D., Chatenay-Lapointe, M., Pan, Y. & Shadel, G. S. Reduced TOR Signaling Extends Chronological Life Span via Increased Respiration and Upregulation of Mitochondrial Gene Expression. Cell Metab. 5, 265–277 (2007).

108. Petsouki, E., Gerakopoulos, V., Gianniou, D. D., Heiss, E. H. & Trougakos, I. P. Involvement of NRF2 and AMPK signaling in aging and progeria: a digest. Redox Biology vol. 85 103782 at 10.1016/j.redox.2025.103782 (2025).

109. Galy, B., Conrad, M. & Muckenthaler, M. Mechanisms controlling cellular and systemic iron homeostasis. Nature Reviews Molecular Cell Biology vol. 25 133–155 at 10.1038/s41580-023-00648-1 (2024).

110. Zhang, D. D. Ironing out the details of ferroptosis. Nat. Cell Biol. 26, 1386–1393 (2024).

111. Lei, G., Zhuang, L. & Gan, B. Targeting ferroptosis as a vulnerability in cancer. Nature Reviews Cancer vol. 22 381–396 at 10.1038/s41568-022-00459-0 (2022).

112. Sun, S., Shen, J., Jiang, J., Wang, F. & Min, J. Targeting ferroptosis opens new avenues for the development of novel therapeutics. Signal Transduction and Targeted Therapy vol. 8 372- at 10.1038/s41392-023-01606-1 (2023).

113. Maus, M. et al. Iron accumulation drives fibrosis, senescence and the senescence-associated secretory phenotype. Nat. Metab. 5, 2111–2130 (2023).

114. Amengual, J. et al. Iron chelation as a new therapeutic approach to prevent senescence and liver fibrosis progression. Cell Death Dis. 15, 680- (2024).

115. Hernandez-Segura, A., Nehme, J. & Demaria, M. Hallmarks of Cellular Senescence. Trends in Cell Biology vol. 28 436–453 at 10.1016/j.tcb.2018.02.001 (2018).

116. Milanovic, M. et al. Senescence-associated reprogramming promotes cancer stemness. Nature 553, 96–100 (2018).

117. Serrano, M. Ageing: Tools to eliminate senescent cells. Nature vol. 545 294–296 at 10.1038/nature22493 (2017).

118. Yamauchi, S. et al. Mitochondrial fatty acid oxidation drives senescence. Sci. Adv. 10, 5887 (2024).

119. Medema, J. P. Escape from senescence boosts tumour growth. Nature vol. 553 37–38 at 10.1038/d41586-017-08652-0 (2018).

120. Wiley, C. D. et al. Mitochondrial dysfunction induces senescence with a distinct secretory phenotype. Cell Metab. 23, 303–314 (2016).

121. Wallace, D. C. Mitochondria and cancer. Nature Reviews Cancer vol. 12 685–698 at 10.1038/nrc3365 (2012).

122. Sabatini, D. M. mTOR and cancer: Insights into a complex relationship. Nature Reviews Cancer vol. 6 729–734 at 10.1038/nrc1974 (2006).

123. Kreuzaler, P., Panina, Y., Segal, J. & Yuneva, M. Adapt and conquer: Metabolic flexibility in cancer growth, invasion and evasion. Molecular Metabolism at 10.1016/j.molmet.2019.08.021 (2020).

124. The importance of aging in cancer research. Nature Aging at 10.1038/s43587-022-00231-x (2022).

125. Vyas, S., Zaganjor, E. & Haigis, M. C. Mitochondria and Cancer. Cell at 10.1016/j.cell.2016.07.002 (2016).

126. Pavlova, N. N., Zhu, J. & Thompson, C. B. The hallmarks of cancer metabolism: Still emerging. Cell Metabolism vol. 34 355–377 at 10.1016/j.cmet.2022.01.007 (2022).

127. Laplante, M. & Sabatini, D. M. MTOR signaling in growth control and disease. Cell vol. 149 274–293 at 10.1016/j.cell.2012.03.017 (2012).

128. van Dijken JP et al. An interlaboratory comparison of physiological and genetic properties of four Saccharomyces cerevisiae strains. Enzyme Microb. Technol. 26, 706–714 (2000).

129. Longtine, M. S. et al. Additional modules for versatile and economical PCR-based gene deletion and modification in Saccharomyces cerevisiae. Yeast 14, 953–61 (1998).

130. Alfatah, M. et al. TORC1 regulates the transcriptional response to glucose and developmental cycle via the Tap42-Sit4-Rrd1/2 pathway in Saccharomyces cerevisiae. BMC Biol. 859793 (2021) doi:10.1186/s12915-021-01030-3.

131. Haque, F., Verma, N. K., Alfatah, M., Bijlani, S. & Bhattacharyya, M. S. Sophorolipid exhibits antifungal activity by ROS mediated endoplasmic reticulum stress and mitochondrial dysfunction pathways in: Candida albicans. RSC Adv. (2019) doi:10.1039/c9ra07599b.

132. Alfatah, M. et al. Metabolism of glucose activates TORC1 through multiple mechanisms in Saccharomyces cerevisiae. Cell Rep. 42, 113205 (2023).

