## Supplemental Figures for "Copper modulates iron-dependent survival through distinct TORC1 and AMPK signaling pathways"

Affiliations:
1. Genome Institute of Singapore (GIS), Agency for Science, Technology and Research (A*STAR), 60 Biopolis Street, Genome #02-01, Singapore 138672, Republic of Singapore.
2. Healthy Longevity Translational Research Program, Yong Loo Lin School of Medicine, National University of Singapore, Singapore, Singapore.
3. Centre for Healthy Longevity, National University Health System, Singapore, Singapore.
4. Bioinformatics Institute (BII), Agency for Science, Technology and Research (A*STAR), 30 Biopolis Street, Matrix #07-01, Singapore 138671, Republic of Singapore.
5. Department of Paediatrics, Yong Loo Lin School of Medicine, National University of Singapore, Singapore 119228, Singapore.
6. Department of Physiology, Yong Loo Lin School of Medicine, National University of Singapore, Singapore 117593, Singapore.
7. Departments of Biochemistry and Physiology, Yong Loo Lin School of Medicine, National University of Singapore, Singapore, Singapore.

#Equal authorship contributions

*To whom the correspondence should be addressed.

 (Mohammad Alfatah)

 (Arshia Naaz)

**Keywords:** Copper; Iron; Redox homeostasis; Cellular survival; TORC1; AMPK; Mitochondria; Nutrient limitation

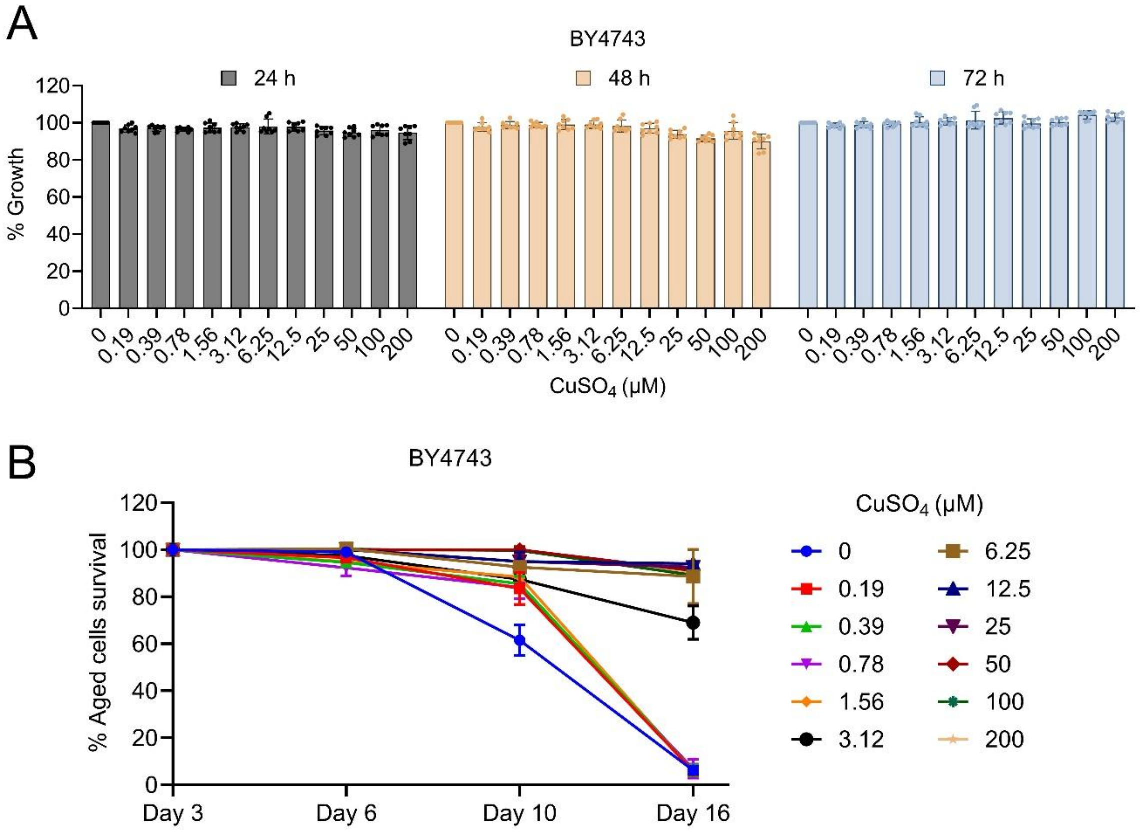


**Figure S1. Copper preserves survival without affecting growth in BY4743 cells, related to Figure 1**

(A) Growth of Saccharomyces cerevisiae BY4743 cells cultured in synthetic defined (SD) medium supplemented with increasing concentrations of $CuSO_4$ (0–200 µM). Cell density was measured at 24, 48, and 72 h by OD600 and normalized to untreated controls.

(B) Chronological survival of BY4743 cells cultured in the presence of $CuSO_4$. Viability was assessed at indicated time points (days 3, 6, 10, and 16) using a liquid outgrowth assay and expressed relative to day 3.

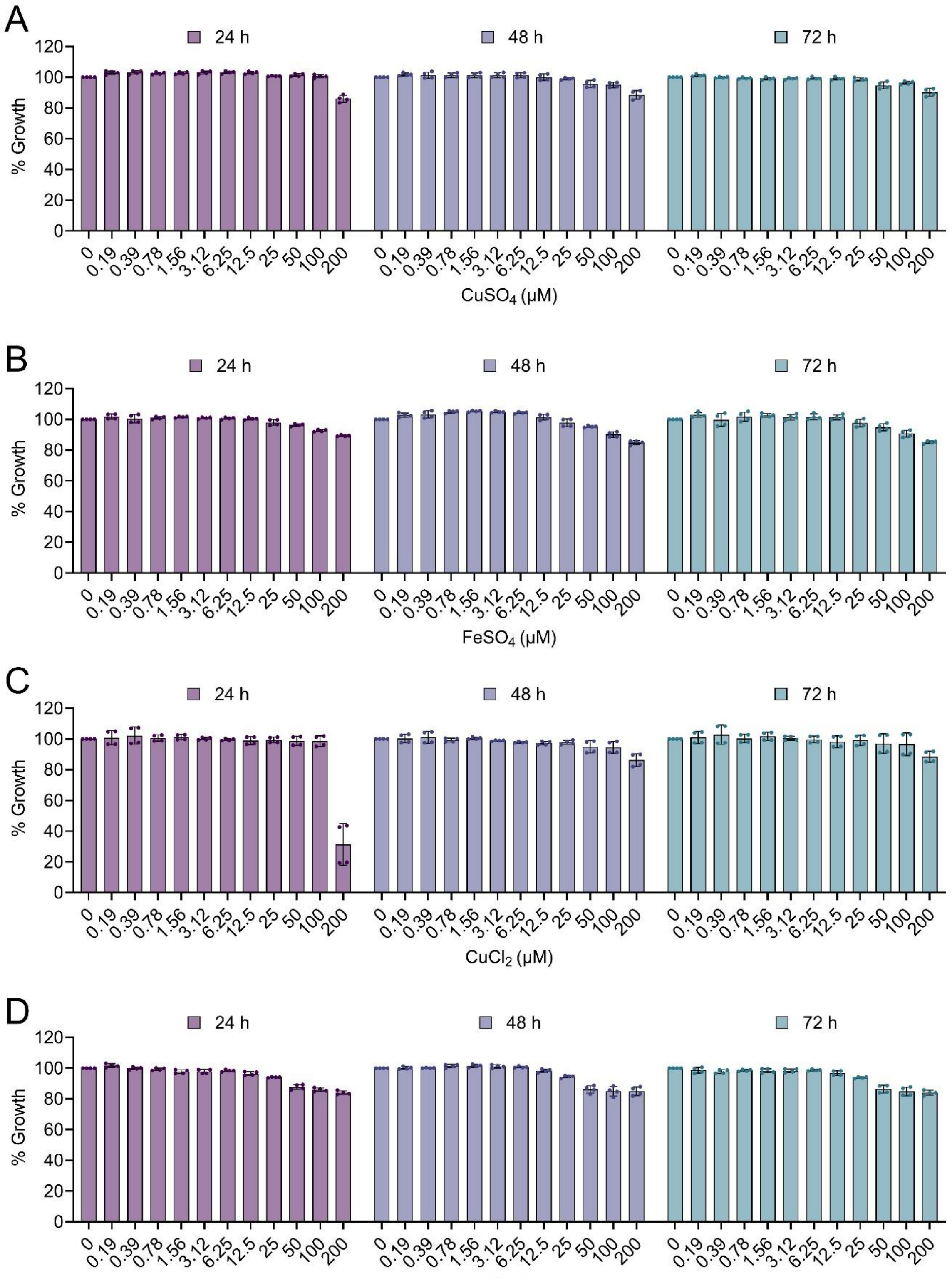


**Figure S2. Copper and iron do not impair proliferative growth, related to Figure 2**

(A) Growth of Saccharomyces cerevisiae CEN.PK cells cultured in synthetic defined (SD) medium supplemented with increasing concentrations of $CuSO_4$ (0–200 μM). Cell density was measured at 24, 48, and 72 h by OD600 and normalized to untreated controls. Data represent mean ± SD (n = 4).

(B) Growth of CEN.PK cells cultured in the presence of $FeSO_4$ (0–200 μM), assessed as in (A). Data represent mean ± SD (n = 4).

(C) Growth of CEN.PK cells cultured in the presence of $CuCl_2$ (0–200 μM), assessed as in (A). Data represent mean ± SD (n = 4).

(D) Growth of CEN.PK cells cultured in the presence of $FeCl_3$ (0–200 μM), assessed as in (A). Data represent mean ± SD (n = 4).

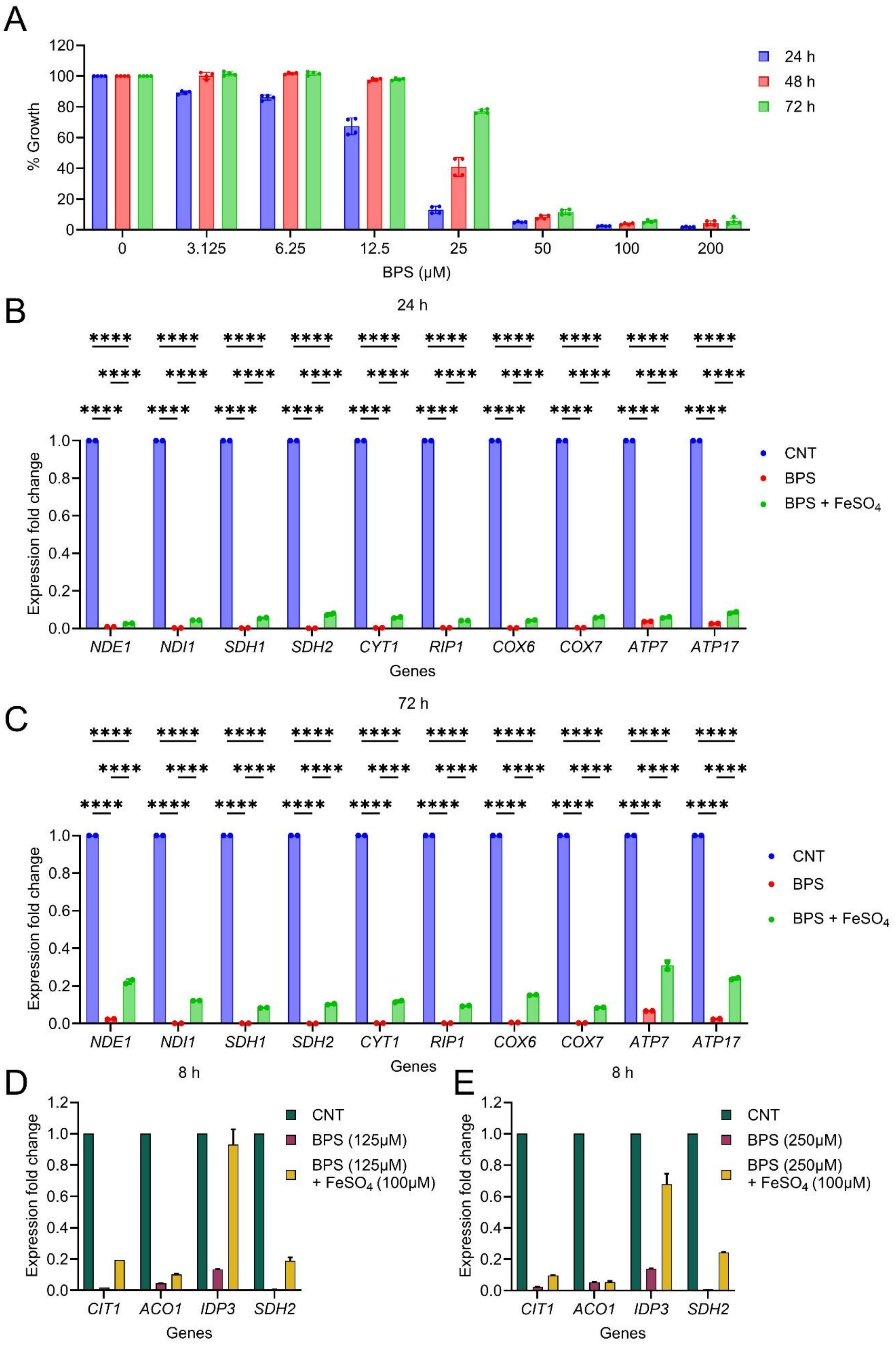
A
% Growth
BPS (μM)
24 h
48 h
72 h
B
24 h
Expression fold change
Genes
CNT
BPS
BPS + FeSO4
NDE1 NDI1 SDH1 SDH2 CYT1 RIP1 COX6 COX7 ATP7 ATP17
C
72 h
D
8 h
CIT1 ACO1 IDP3 SDH2
BPS (125μM)
BPS (125μM) + FeSO4 (100μM)
E
BPS (250μM)
BPS (250μM) + FeSO4 (100μM)

**Figure S3. Iron supplementation restores mitochondrial gene expression under iron chelation, related to Figure 3**

(A) Growth of *Saccharomyces cerevisiae* CEN.PK cells cultured in synthetic defined (SD) medium with increasing concentrations of the iron chelator BPS (0–200 μM). Cell density was measured at 24, 48, and 72 h by OD600 and normalized to untreated controls. Data represent mean ± SD (n = 4).

(B) Relative expression of mitochondrial electron transport chain–related genes (*NDE1, NDI1, SDH1, SDH2, CYT1, RIP1, COX6, COX7, ATP7,* and *ATP17*) in control (CNT), BPS-treated, and BPS + $FeSO_4$-treated cells at 24 h. Gene expression was measured by quantitative RT–PCR, normalized to *ACT1*, and expressed relative to control. Data represent mean ± SD (n = 2). Statistical significance was assessed by two-way ANOVA with Tukey's multiple comparisons test; ****P < 0.0001.

(C) Relative gene expression as in (B), measured at 72 h. Data represent mean ± SD (n = 2). Tukey's multiple comparisons test; ****P < 0.0001.

(D) Relative expression of tricarboxylic acid (TCA) cycle–related genes (*CIT1, ACO1, IDP3,* and *SDH2*) in control (CNT), BPS (125 μM), and BPS + $FeSO_4$-treated cells at 8 h. Gene expression was normalized to *ACT1* and expressed relative to control. Data represent mean ± SD (n = 2).

(E) Relative gene expression of TCA cycle–related genes as in (D), measured under BPS (250 μM) conditions at 8 h. Data represent mean ± SD (n = 2).

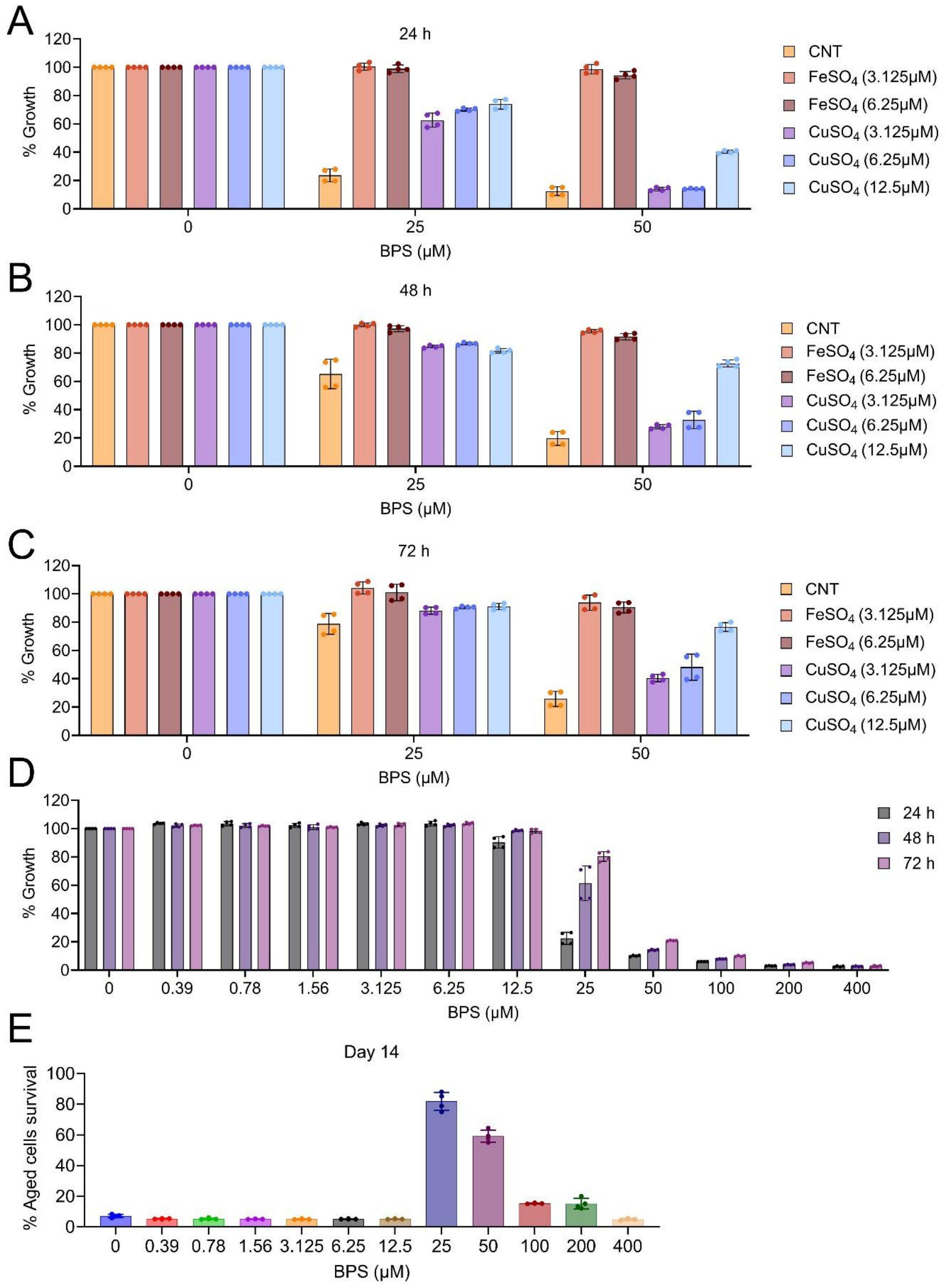


**Figure S4. Iron supplementation more effectively restores growth than copper under iron chelation, related to Figure 4**

(A–C) Growth of *Saccharomyces cerevisiae* CEN.PK cells cultured in synthetic defined (SD) medium with BPS (0, 25, and 50 μM) in the presence of $FeSO_4$ (3.125 and 6.25 μM) or $CuSO_4$

(3.125, 6.25, and 12.5 µM). Cell density was measured at 24 h (A), 48 h (B), and 72 h (C) by OD600 and normalized to untreated controls (CNT). Data represent mean ± SD (n = 4).

(D) Growth of CEN.PK cells cultured with increasing concentrations of BPS (0–400 µM). Cell density was measured at 24, 48, and 72 h by OD600 and normalized to untreated controls. Data represent mean ± SD (n = 4).

(E) Chronological survival of CEN.PK cells cultured with increasing concentrations of BPS (0–400 µM). Viability was assessed at day 14 using a liquid outgrowth assay and expressed relative to day 3. Data represent mean ± SD (n = 4).

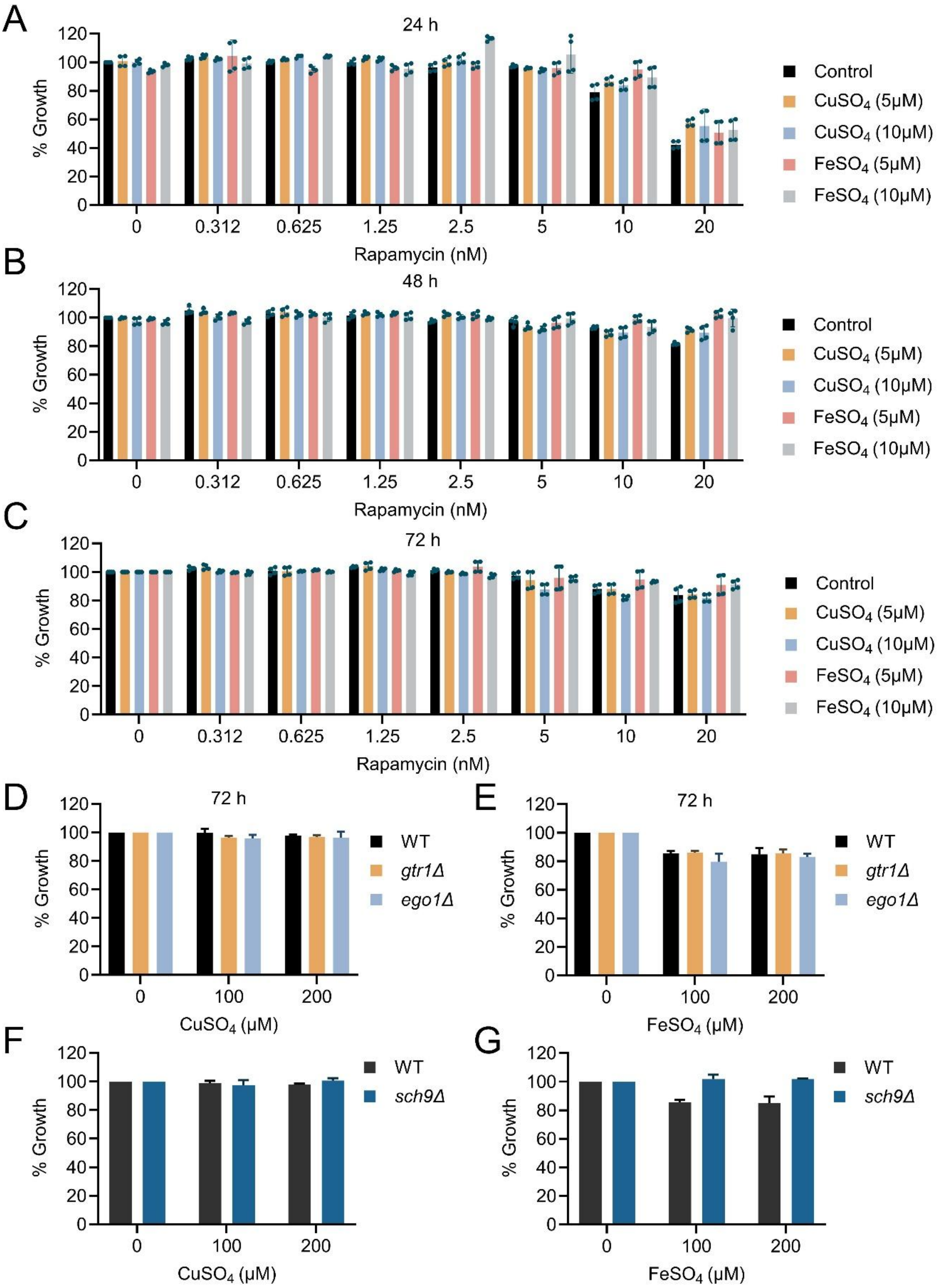


**Figure S5. Copper and iron do not substantially alter proliferative growth under TORC1 perturbation or genetic backgrounds, related to Figure 5**

(A–C) Growth of *Saccharomyces cerevisiae* CEN.PK cells cultured in synthetic defined (SD) medium with increasing concentrations of rapamycin (0–20 nM) in the presence or absence of $CuSO_4$ (5 and 10 μM) or $FeSO_4$ (5 and 10 μM). Cell density was measured at 24 h (A), 48 h (B), and 72 h (C) by OD600 and normalized to untreated controls. Data represent mean ± SD (n = 4).

(D–E) Growth of wild-type (WT), *gtr1Δ*, and *ego1Δ* strains cultured in the presence of $CuSO_4$ (0–200 μM) (D) or $FeSO_4$ (0–200 μM) (E). Cell density was measured at 72 h by OD600 and normalized to WT control. Data represent mean ± SD (n = 2).

(F–G) Growth of WT and *sch9Δ* strains cultured in the presence of $CuSO_4$ (0–200 μM) (F) or $FeSO_4$ (0–200 μM) (G). Cell density was measured by OD600 and normalized to WT control. Data represent mean ± SD (n = 2).

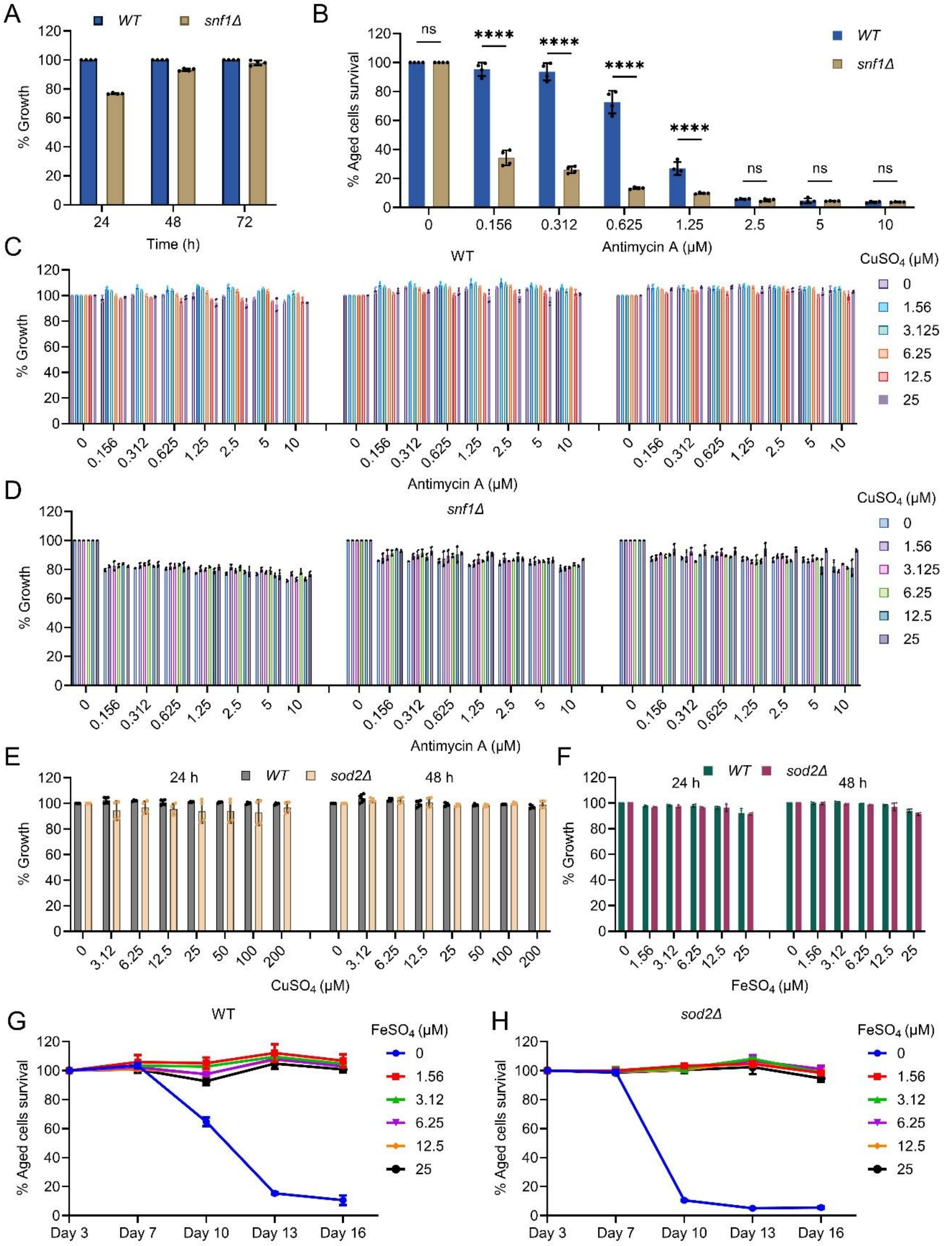


**Figure S6. AMPK/Snf1 and antioxidant capacity influence stress responses without affecting proliferative growth, related to Figure 6**

(A) Growth of wild-type (WT) and *snf1Δ* strains cultured in synthetic defined (SD) medium. Cell density was measured at 24, 48, and 72 h by OD600 and normalized to WT control. Data represent mean ± SD (n = 4).

(B) Chronological survival of WT and *snf1Δ* cells cultured with increasing concentrations of antimycin A (0–10 μM). Viability was assessed using a liquid outgrowth assay and expressed relative to day 3. Data represent mean ± SD (n = 4). Statistical significance was assessed by two-way ANOVA with Šídák's multiple comparisons test; ****$P < 0.0001$; ns, not significant.

(C–D) Growth of WT (C) and *snf1Δ* (D) cells cultured with increasing concentrations of antimycin A (0–10 μM) in the presence of $CuSO_4$ (0–25 μM). Cell density was measured by OD600 and normalized to untreated controls. Data represent mean ± SD (n = 4).

(E–F) Growth of WT and *sod2Δ* strains cultured with increasing concentrations of $CuSO_4$ (0–200 μM) (E) or $FeSO_4$ (0–25 μM) (F). Cell density was measured at 24 and 48 h by OD600 and normalized to WT control. Data represent mean ± SD (n = 4).

(G–H) Chronological survival of WT (G) and *sod2Δ* (H) strains cultured with increasing concentrations of $FeSO_4$ (0–25 μM). Viability was assessed at indicated time points and expressed relative to day 3. Data represent mean ± SD (n = 2).

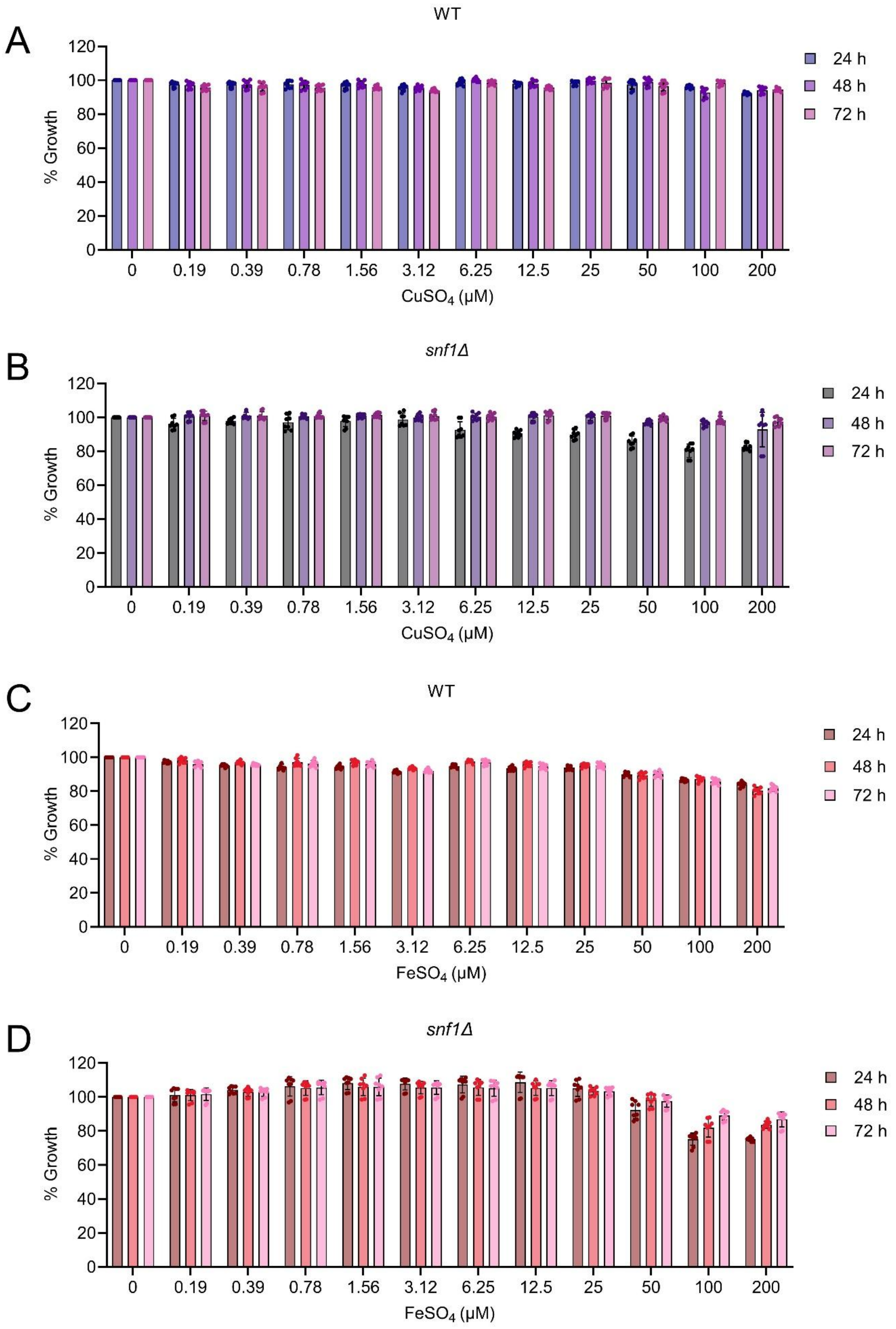
A
WT
% Growth
CuSO4 (μM)
0
0.19
0.39
0.78
1.56
3.12
6.25
12.5
25
50
100
200
24 h
48 h
72 h
B
snf1Δ
% Growth
CuSO4 (μM)
C
WT
% Growth
FeSO4 (μM)
D
snf1Δ
% Growth
FeSO4 (μM)

**Figure S7. Copper and iron do not impair proliferative growth in WT and *snf1Δ* strains, , related to Figure 7**

(A–B) Growth of wild-type (WT) (A) and *snf1Δ* (B) cells cultured in synthetic defined (SD) medium supplemented with increasing concentrations of $CuSO_4$ (0–200 μM). Cell density was measured at 24, 48, and 72 h by OD600 and normalized to untreated controls. Data represent mean ± SD (n = 8).

(C–D) Growth of WT (C) and *snf1Δ* (D) cells cultured in SD medium supplemented with increasing concentrations of $FeSO_4$ (0–200 μM). Cell density was measured at 24, 48, and 72 h by OD600 and normalized to untreated controls. Data represent mean ± SD (n = 8).
